# Retro-miRs: Novel and functional miRNAs originated from mRNA retrotransposition

**DOI:** 10.1101/2023.02.24.529967

**Authors:** Rafael L. Mercuri, Helena B. Conceição, Gabriela D. A. Guardia, Gabriel Goldstein, Maria D. Vibranovski, Ludwig C. Hinske, Pedro A F Galante

## Abstract

**Background:** Reverse transcribed gene copies, or retrocopies, have emerged as a major source of evolutionary novelties. MicroRNAs (miRNAs) are small, highly conserved RNAs molecules among species that serve as key post-transcriptional regulators of gene expression. The birth and subsequent evolution of miRNAs have been addressed, but not fully.

**Results:** In this study, we carried out a comprehensive investigation of miRNAs origination through retroduplicated mRNA sequences (retrocopies). We identified 17 retroduplicated miRNAs (retro-miRs) that emerged from mRNAs retrocopies. Four of these retro-miRs had *de novo* origination within retrocopied sequences, while 13 retro-miRNAs were located within exon regions and were duplicated along with their host mRNAs. We found that retro-miRs are primates specific, including 5 retro-miRs conserved among all primates and two human-specific retro-miRs. All of the retro-miRs were expressed and had predicted and experimentally validated target genes, with the exception of miR-10527. Notably, the target genes of retro-miRs are involved in key biological processes, such as metabolic processes, cell signaling and regulation of neurotransmitters in the central nervous system. Additionally, we found that these retro-miRs have a potential oncogenic role in cancer, targeting key cancer genes and being overexpressed in several cancer types, including Liver Hepatocellular Carcinoma and Stomach Adenocarcinoma.

**Conclusion:** Our findings demonstrate that mRNAs retrotransposition is a key mechanism for the generation of novel miRNAs (retro-miRs) in primates. These retro-miRs are expressed, conserved, have target genes with important cellular functions, and play roles in cancer.

## INTRODUCTION

MicroRNAs (miRNAs) are small non-protein coding RNAs (approximately 22 bp) with a key function of regulating gene expression at the post-transcriptional level [1]. Advances in understanding miRNA biology have already revealed hundreds of miRNAs genes widespread throughout the animal kingdom [2]. The biogenesis of miRNAs involves several coordinated processing steps, resulting in a mature miRNA that is incorporated in the RNA-induced silencing complex (RISC) [3]. miRNAs act in gene regulation mechanisms by targeting 3’ UTR protein coding genes leading to translational repression or mRNA degradation [4]. Additionally, it has been shown that miRNAs act as gene regulators very early on in the evolution of animals and have complex evolutionary patterns, different from other genetic sequences [5].

Remarkably, it has been shown that the genome incorporation of novel miRNAs is a pivotal and instrumental step in the evolution of organismal complexity [5]. It is clear that the expression of miRNAs modulates the transcript variability of their target coding gene, fine tuning the protein molecules that the coding gene produces, and therefore, conferring robustness to cellular pathways and gene networks. Consequently, this gene expression robustness may have contributed to a decrease of phenotypic variation, maintaining invariant phenotype in the face of endogenous and exogenous perturbations. This would increase heritability of species-specific traits [6].

For the emergence of new sequences of miRNAs, it is a fundamental prerequisite that the novel miRNA gene is transcribed from a genomic locus prone to produce an RNA fold recognizable by the miRNA processing machinery [7]. Essentially, there are two distincts molecular mechanisms capable of originating a novel miRNA: i) the duplication of a pre-existing miRNA, followed by sub or neofunctionalization processes [8]; and ii) de novo origination of miRNAs [9]. While the former is majorly mediated by a local or full genome duplication [10], the latter (*de novo*) miRNA origination has a bias to occur in transcribed regions, frequently from introns [11,12]) or mediated by transposable elements [13] in duplication of non-coding RNAs [14–16].

One overlooked source for novel miRNAs are copies of processed mRNA, also known as retrocopies. Retrocopies are mRNA copies (of protein-coding genes) that have been reverse transcribed to cDNA sequence and re-inserted into the genome in a process known as retrotransposition [17]. Until recently, retrocopies have been referred to as processed pseudogenes, but an increasing body of evidence suggests that a large fraction of retrocopies are functional [18–20]. Now, it is clear that retrocopies are a major source of genetic novelties by creating novel genes [21], regulatory regions [22] and other non-coding genes, including miRNAs [15]. Specifically, this last work showed that two miRNAs (hsa-miR-220 and hsa-miR-492) lie within retrocopies of protein coding genes and suggest that these retroduplicates are good candidates as “miRNA incubators’’. Surprisingly, almost two decades later no further investigations have been performed regarding this issue.

In this study, we comprehensively investigate the contribution of retrocopies as a source of novel miRNAs (retrocopy derived miRNAs, retro-miRs) in the human genome. To accomplish this investigation, we used an extensive range of databases concerning genomic annotations, sequence conservation, expression in health and cancer tissues, information about miRNA target genes and a complete set of bioinformatics tools. In summary, we unveiled 17 primate specific retro-miRs, investigated their conservation, expression, gene targets and putative function in cancer.

## 2 RESULTS

### 2.1 Finding miRNAs originated by mRNA retrotransposition events

To identify and investigate miRNAs originated from mRNA retrocopies (retro-miRs), we developed a set of local pipelines and surveyed several databases, Figure 1A. Briefly, we constructed and used a set of computational algorithms to assess and integrate information from three databases: miRBase, the reference database to several miRNAs’ information [23]; miRIAD, a database containing annotation and further data of intragenic miRNAs and their host genes [24,25]; and RCPedia, a database of retrocopies present in humans and other species [26]). Further information about the investigated genes (retrocopies and miRNAs), such as their conservation, expression, miRNAs target genes, and functional information were retrieved from other databases (e.g., TargetScan [27], miRTarBase [28], FANTOM phase 5, and TCGA) and locally processed, Figure 1A. See Methodology for detail.

**Figure 1:**
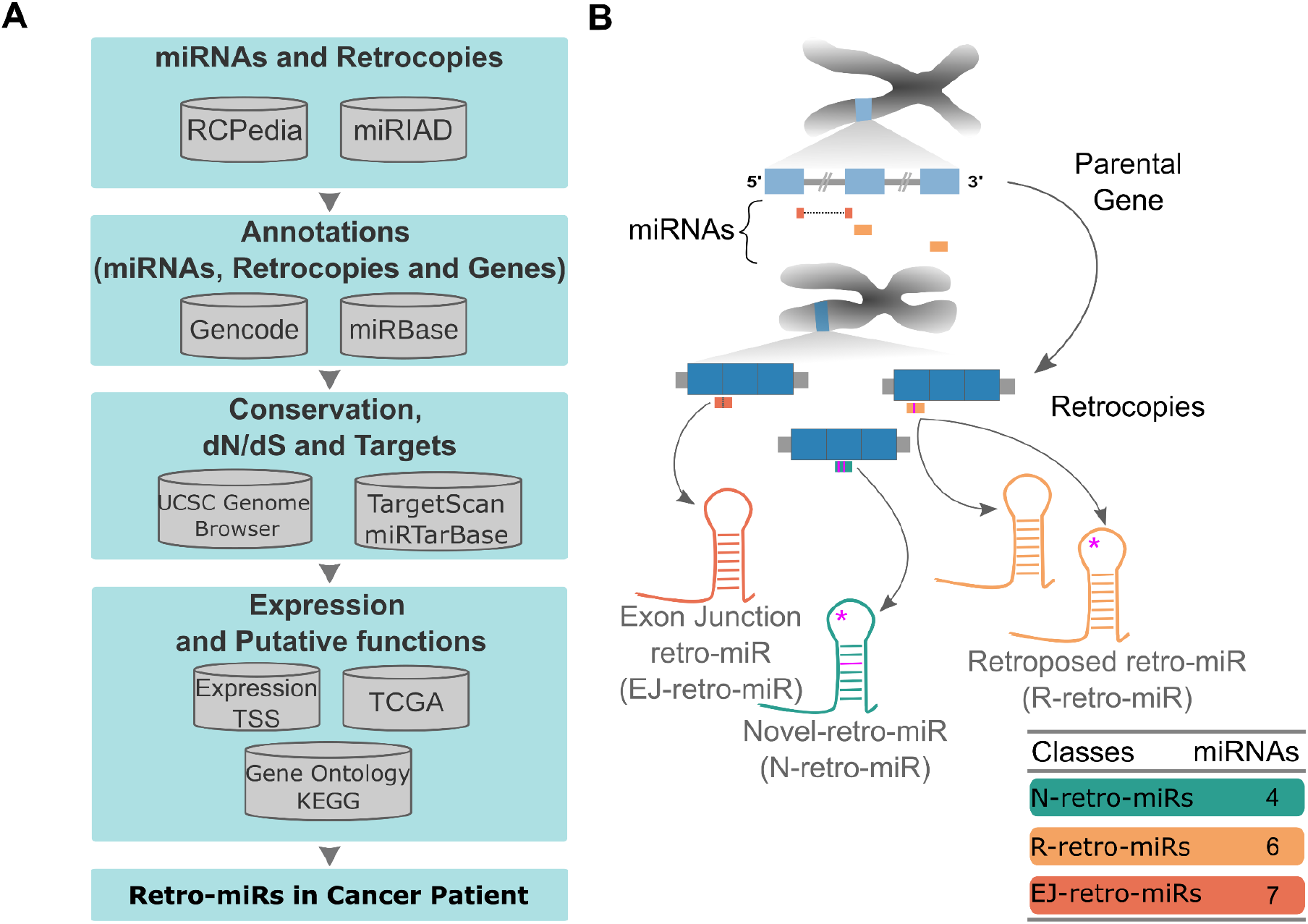
Finding miRNAs originated by retrotransposition of protein coding genes. **A)** A summary of databases, datasets, and strategy used to identify retrocopy derived miRNAs (retro-mirs). **B)** A schematic representation of a parental gene, exonic miRNAs, retrocopies and retro-miRs. All three classes of retro-miRs, exon-junction retro-miRs (EJR-miR, in red), novel retro-miRs (NR-miR, in green), and retroposed retro-miRs (RR-miR, in orange) are represented. Tiny purple bars in retro-miRs representations and harpins represent mismatches between them and their respective region in the parental gene.

By using this strategy, we found 17 retrocopy derived miRNAs (retro-miRs) laying into retrocopied sequences. These miRNAs were grouped in three distinct classes: Retroposed (R-retro-miRs), Exon Junction EJ-retro-miRs and Novel retro-miRs (N-retro-miRs), Figure 1B. Six R-retro-miRs (35.3%; mir-4444-2; mir-1244-2; mir-1244-3; mir-1244-4; mir-1244-5; mir-1244-6) are retroduplications of exonic miRNAs already present in their parental gene sequences (Figure 1B, Supplementary Figure S1). These include four R-retro-miRs fully identical to their parental miRNAs and two others that have variation in their precursor stem, including one retro-miR with variation in the mature region (out seed region). We found seven EJ-retro-miRs (41.2%; mir-492, mir-622, mir-4426, mir-4426-1, mir-3654, mir-3654-1, and mir-10527), which are also in exonic region of their retrocopies’ parental genes, but spanning two exons (Figure 1B, Supplementary Figure S2). Curiously, none of these miRNAs have ever been annotated in the parental gene region, likely because they are splitted by an intronic sequence. The last retro-miRs class, N-retro-miR, comprise four miRNAs (23.5%; mir-4788, mir-7161, mir-4468, and mir-572) which had a de novo origination in the retrocopied sequences due to the occurrence of mutations along evolution (Figure 1B, Supplementary Figure S3). Upon checking the equivalent region in the parental gene, no evidence of a stem loop, i.e., a putative miRNA was found (Supplementary Figure S4).

Next, we further investigated characteristics of these retrocopies containing retro-miRs. First, we found that the six R-retro-miRs present in both retrocopy and parental genes, were originated in six distinct retroduplication events, but from two parental genes only: PTMA (5 retrocopies with retro-miRs) and HNRNPA3 (one retrocopy), Supplementary Table S1. Interestingly, these genes are highly retroduplicated in the human genome. PTMA presents 13 retrocopies and HNRNPA3 presents 17 retrocopies in total (Supplementary Table S2). On the other hand, 5 distinct parental genes originated the 7 retrocopies with 7 EJ-retro-miRs and four distinct parental genes that originated four retrocopies, each with a N-retro-miRs, Supplementary Table S1 and S2.

These previous results led us to investigate events of chromosomal duplication of genes containing exonic miRNAs, in addition to the retro-miRs we identified. Briefly, we found that among the 19,768 protein coding genes, 144 protein coding genes contain exonic miRNAs, where 17 of these protein coding genes were duplicated by retrotransposition and 24 by the DNA-mediated duplication mechanism. Interestingly, only two of the DNA-based duplications also have miRNAs in their duplicates (Supplementary Table S3). All together, these results suggest that retroduplication is a frequent pathway for novel miRNA’s origin.

### 2.2 Characteristics and conservation of retro-miRs events

To trace the evolutionary origin of each retrocopy and their retro-miRs, we used a homology-based approach (see Methods) to query their conservation, Figure 2A. First, this analysis revealed that all of these retrocopies and their miRNAs originated in the primate lineage (Supplementary Table S4) and, in agreement with literature of retrocopies, are primate specific [19]. By correlating the age of retro-miRs and their classes, we observed that the oldest events (conserved in all primates), are enriched with *de novo* N-retro-miRs. While EJ-retro-miRs are spread in all ages and R-retro-miRs are only in great apes and up, Figure 2A.

These results prompted us to check whether retrocopies containing retro-miRs have evolved under a functional constraint (purifying or positive selection) or neutrality. To assess this, we used retrocopies’ predicted coding region to quantify the ratio of the number of nonsynonymous substitution per nonsynonymous site to the number of synonymous substitution per synonymous site (dN/dS). We found three retrocopies clearly under neutral selection (dN/dS ∼ 1; retrocopies containing retro-mir-4444-2; retro-mir-1244-2 and retro-mir-3654-1) and 12 retrocopies with a dN/dS ratio different from 1 (potentially under functional constraint), Figure 2D. However, only for three of these retrocopies we confirmed a dN/dS significantly different than 1 (p-value <= 0.05, likelihood ratio test), being two retrocopies under purifying selection (retrocopies containing retro-mir-1244-6 and retro-mir-7161) and one retrocopy (containing retro-mir-1244-5) under positive selection, Figure 2D, Supplementary Table S5.

**Figure 2.**
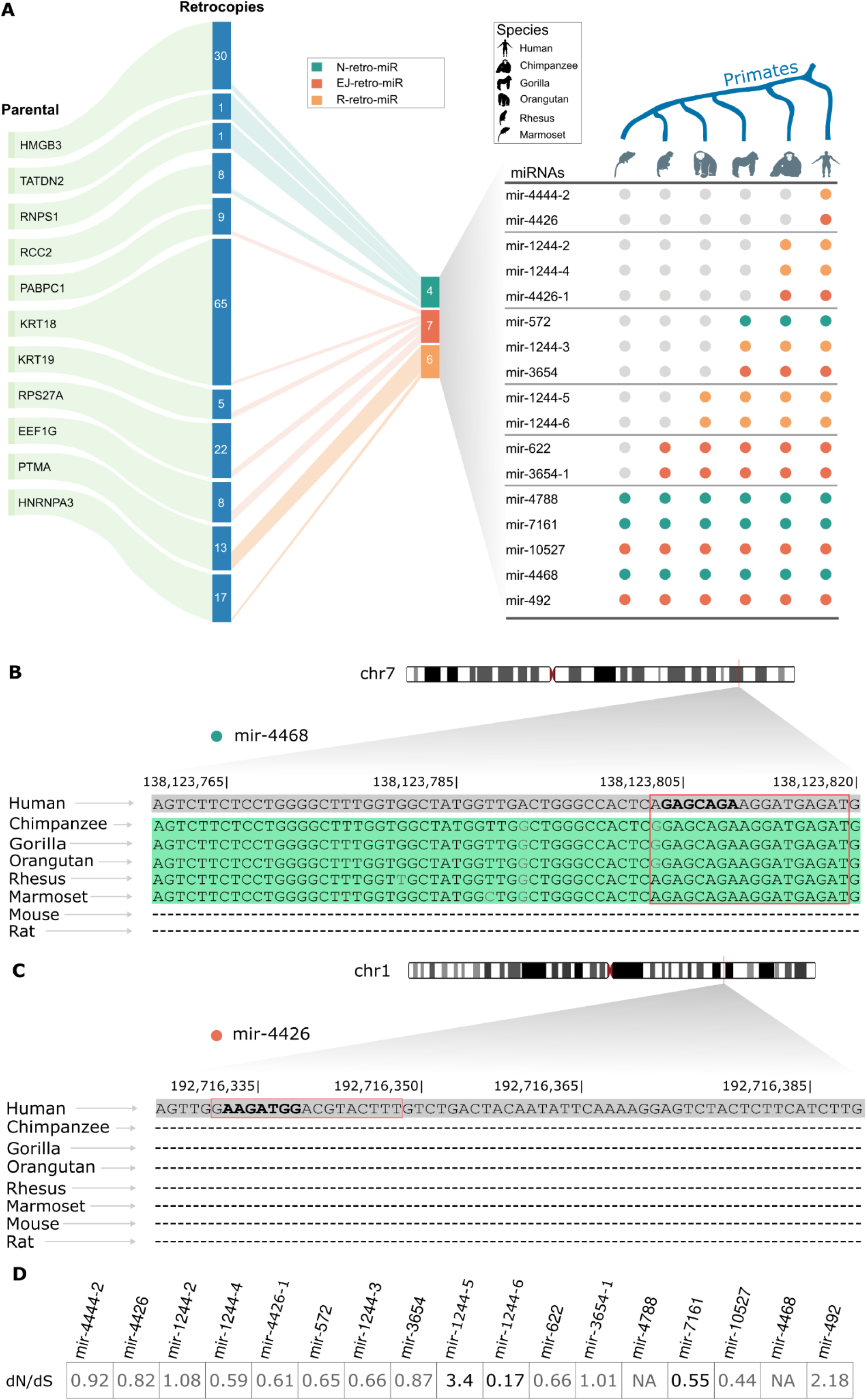
Retro-miRNAs originated in primate lineage and have evolved under a functional constraint. **A)** We identified 17 retro-miRs that originated from 11 parental genes (left side). On the right side, we show the conservation of these retro-miRs in human, chimpanzee, gorilla, orangutan, rhesus, and marmoset genomes. Gray markings indicate the absence of the sequence in that organism. **B)** An alignment between primate, mouse, and rat genomes of a retro-miR (mir-4468, a retro-miR conserved among all primates) sequence. The regions conserved are highlighted in green (matches in black; mismatches in light gray), and “-” indicates a lack of conservation. The red boxes represent the mature mRNA sequence, and the bold sequences are the miRNA seed region (2nd to 8th nt). **C)** Alignment of a human-specific retro-miR (mir-4426). Lack of conservation is presented by “-”. The red boxes represent the mature sequence, and the bold sequences are the miRNA seed region (2nd to 8th nt). **D)** Retro-miRs in retrocopies with a dN/dS significantly different than 1 (p-value <= 0.05, likelihood ratio test) under purifying (dN/dS < 1) or positive (dN/dS > 1) selection are highlighted in black, while those with non-significant (or non-evaluated “NA”) dN/dS values are shown in gray.

All together, these results indicated that retro-miRs originated along primate evolution in distinct waves. Addictionaly, considering the recent insertion of these retrocopies and retro-miRs in the evolutionary history of humans, we see a tendency for purifying evolutionary pressure on them.

### 2.3 Expression of retro-miRs in normal tissues

Having dissected the genomic features of retro-miRs, we moved forward to identify their transcription and target genes, a fundamental feature of functional miRNAs. First, we sought for evidence of retro-miR transcription using short RNA-sequencing [29] data and CAGE data [30]. We found that all retro-miRs (except retro-miR-10527) showed expression in normal samples (404 individuals, 32 tissues) and/or cell lines (48 samples of stem cells), Figure 3A. Notably, we observed a distinct expression pattern among the retro-miRs in the different tissues and cell lines (stem cells). Interestingly, a novel retro-miR (mir-4788) is highly expressed (the highest median expression among all retro-miRs expressed) in normal samples. Conversely, in stem cells this retro-miR (mir-4788) has the lowest (median) expression.

**Figure 3.**
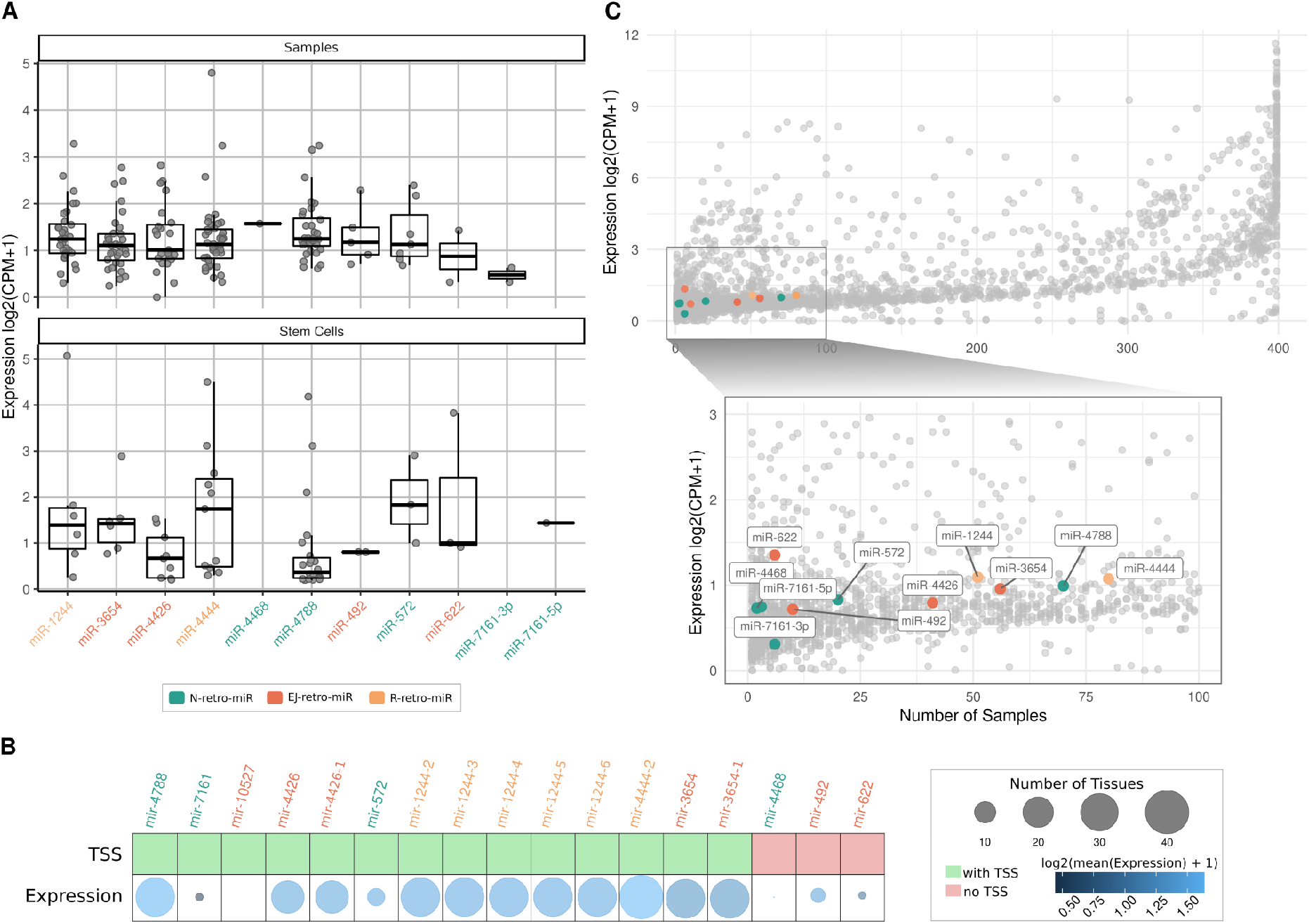
Retro-miRNAs are expressed in normal tissues and cell lines. **A)** Expression (median per tissue) of retro-miRs in normal samples (404 individuals, 32 tissues) and cell lines (48 samples of stem cells). **B)** A table showing the presence (in green) or absence (in red) of a transcription start site (TSS) upstream of the retro-miR, and a bubble plot representing the intensity of expression of retro-miRs, as well as the number of tissues in which they were expressed. **C)** Expression level per number of tissues of all known human miRNAs. Retro-miRs are marked in colors, while all other miRNAs are represented by gray dots. Only expressed retro-miRs are shown in A and C.

Next, to further support the true expression of these loci containing retro-miRs, we evaluated the presence of a transcription start site (TSS) near (up to 4K bp) upstream their 5’ end using CAGE sequencing. This data confirmed TSSs for 82% (14/17) retro-miRs, Figure 3B. In agreement with RNAseq data, retro-miRs with a defined TSS have, in general, a higher expression and are more broadly expressed in different tissues than retro-miRs without a TSS, Figure 3B.

Next, we investigate the expression of retro-miRs in comparison to all other annotated miRNAs (Supplementary Table S6). Figure 3C shows that retro-miRs are in the middle of the first quartile in terms of expression levels and number of tissues expressed. Together, we observe that all retro-miRs have a similar expression, which is expected for genes (miRNAs) that originated recently in evolution. Additionally, N-retro-miRs are among the miRNAs expressed in fewer tissues, a pattern also expected for novel genes (miRNAs).

In a further investigation, we sought to identify the set of genes targeted by retro-miRs, a *sine qua non* feature of functional miRNAs [27]. Notably, we identify target genes for all retro-miRs using TargetScan (7mer-m8 and 8mer), targetscan (8mer) and also experimentally validated targets, Figure 4A (Supplementary Table S7). Only one retro-miR (mir-10527) has no experimentally validated targets, in accordance with our previous result (Figure 3, legend), where we report no expression for mir-10527. Next, in order to globally evaluate the functions of these retro-miRs’ targets, we performed a Gene Ontology investigation of their biological processes. We found that these target genes carry out function in key cellular processes, including regulation of transcription, DNA replication, cell proliferation and differentiation, DNA binding, among others, Figure 4B (p-value < 0.05; fold enrichment > 1.5). Next, we separately checked each set of targets from all retro-miRs (Supplementary Figure S5). Again, two very interesting sets of target genes emerged. The two novel retro-miRs (mir-4788 and mir-572) have gene targets enriched to neural biological processes, specially genes involved in regulation of neurotransmitter and synapses, Figure 4C (Supplementary Figure S5). In addition, mir-4788 is conserved from Humans to Marmoset, and mir-572 is conserved from Humans to Gorilla. This finding could be implicated with the phenomenon of species specific traits heritability known to miRNA.

**Figure 4.**
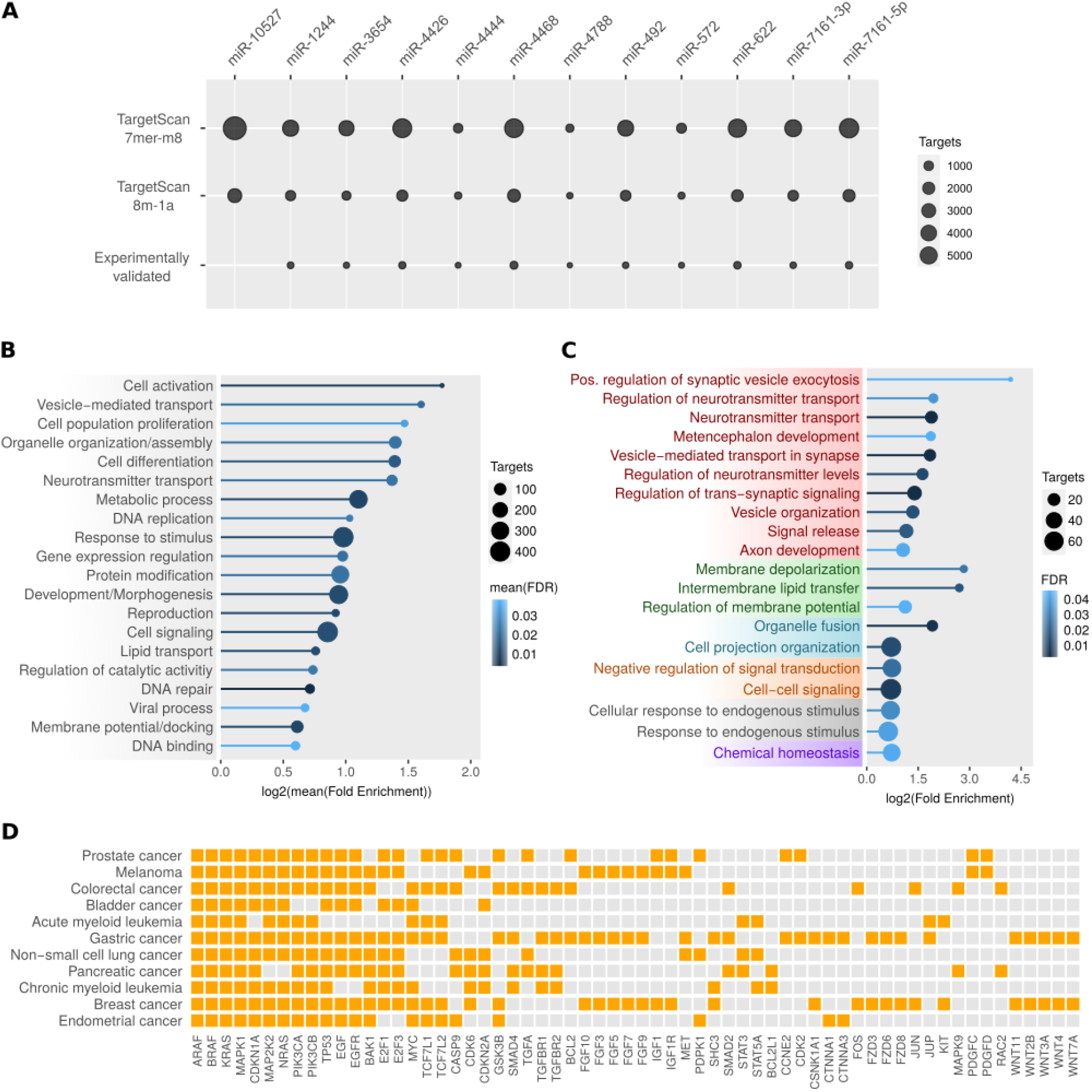
Retro-miRs have target genes carrying out key cell functions, including regulation in neural tissues, and with oncogenic roles. A) Number of predicted (7mer-m8 and 8m-1a) and experimentally validate target genes for retro-miRs. B) Gene ontology enrichment (Biological Processes) analysis of gene targets from retro-miRs. C) Gene ontology enrichment (Biological Processes) analysis of target genes from mir-572 and mir-4788. D) Cancer-related KEGG pathways enriched in target genes of retro-miRs. Only enriched (> 1.5) and significant processes and pathways (FDR < 0.05) are shown.

We also assessed whether retro-miRs’ targets were enriched in particular KEGG pathways. Remarkably, we found enrichment of several cancer-specific pathways, including breast, colorectal, gastric, pancreatic, prostate and lung cancers (Figure 4D), in addition to more general cancer-related pathways, such as P53, ErbB and MAPK signaling, as well as platinum drug and EGFR tyrosine kinase inhibitor resistance (Supplementary Table S8). In this analysis, several known cancer genes emerged as retro-miRs’ targets, including BRAF, KRAS, EGFR and TP53.

All together, the expression and gene targets investigation revealed that retro-miRs (except mir-10527) have reliable expression and validated gene targets, two fundamental characteristics of functional miRNAs. We also observed that retro-miRs have target genes in fundamental cell processes (e. g., cell growth, regulation of transcription and post-transcriptional regulation) and that two novel retro-miRs (mir-572 and mir-4788) regulate genes majorly related to neuronal functions. Additionally, we observed that these retro-miRs have key cancer genes among their targets, suggesting that they play oncogenic roles in cancer tissues.

### 2.4 Retro-miRs are highly or majorly transcribed in cancer tissues

Our previous results (Figure 4D) prompted us to study the expression of these retro-miRs in cancer tissues. By using the short-RNA sequencing data available in The Cancer Genome Atlas (TCGA; [31]), we sought for expression of mature retro-miRs in 10 solid tumors. Their normal counterpart tissues were used as control, despite their relatively limited number of normal samples. Overall, this investigation revealed that retro-miRs have a distinct and commonly high expression in cancer (Figure 5). More specifically, miR-10527-5p, miR-3654, miR-4444, miR-4788, and miR-1244 are highly expressed in cancer, but are also expressed in several normal tissues (Figure 5). Curiously, miR-10527-5p is highly expressed in tumors, despite not being expressed in normal samples or cell lines (Figure 3). Remarkably, five retro-miRs emerged as putative tumor biomarkers because they present a very low (or absent expression) in normal samples and a consistent expression in cancer. Notably, two retro-miRs (miR-622 and miR-492) are exclusively expressed in seven cancer types each, BRCA (miR-622), COAD (miR-622 and miR-492), LUAD (miR-622), PAAD (miR-622 and miR-492), PRAD (miR-622 and miR-492), STAD (miR-492), and THCA (miR-492), Figure 5. Two matures from the same precursor (mir-7161), miR-7161-3p and miR-7161-5p, also have an exclusive expression in six cancer types, BRCA (miR-7161-3p and miR-7161-5p), COAD (miR-7161-3p), LUAD (miR-7161-3p), LIHC (miR-7161-5p), PAAD (miR-7161-3p and miR-7161-5p), and PRAD (miR-7161-3p). Finally, miR-4426 is exclusively expressed in BRCA, COAD, KIRC, and PAAD.

**Figure 5.**
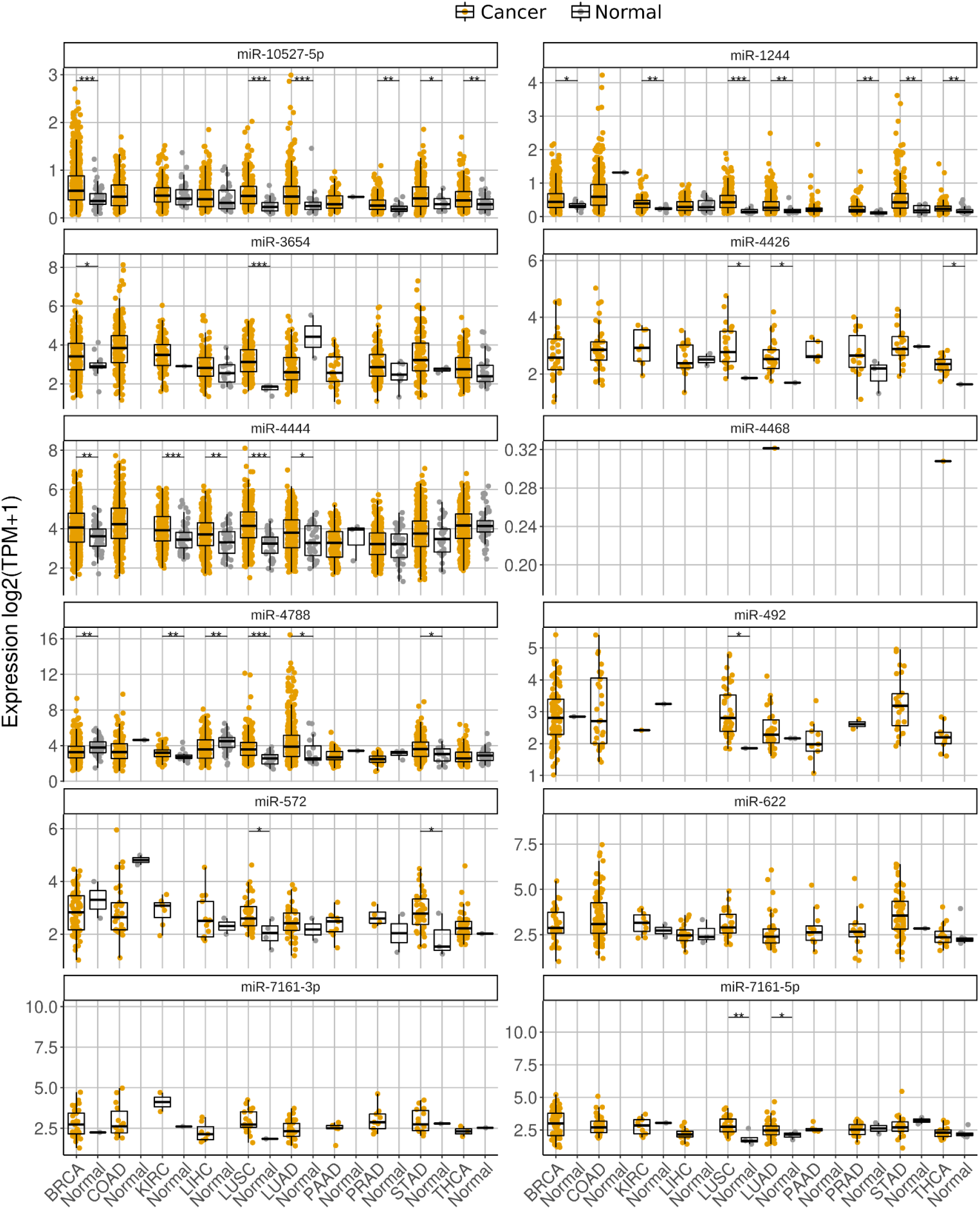
Retro-miRNAs are highly expressed in Cancer and some retro-miRs present a cancer specific expression. Expression level of (mature) retro-miRs in different types of tumors (BRCA = Breast cancer, COAD = Colon adenocarcinoma, KIRC = Kidney renal clear cell carcinoma, LIHC = Liver hepatocellular carcinoma, LUSC = Lung squamous cell carcinoma, LUAD = Lung adenocarcinoma, PAAD = Pancreatic adenocarcinoma, PRAD = Prostate adenocarcinoma, STAD = Stomach adenocarcinoma, THCA = Thyroid carcinoma) and their respective normal samples. Wilcoxon test Tumor x Healthy samples: p-value <= 0.05 and >= 0.01 (*); <0.01 and >= 0.0001 (**); <0.0001 (***)

All together, these results indicate that retro-miRs are highly expressed in cancer, with some retro-miRs presenting a cancer-specific expression pattern and a potential as tumor biomarkers to be further studied.

### 2.5 Overall Survival based on groups of retro-miRs in TCGA samples

Having confirmed the high and distinct expression of retro-miRs in cancer, we sought to check whether these miRNAs have prognostic values to overall survival in cancer patients. To assess this, we used Reboot [32], an algorithm to find gene signatures (including miRNAs) associated with cancer patient prognosis. Using this strategy in approximately 8 thousand samples from 10 cancer types presenting overall survival (OS), we found two signatures with prognostic values, a signature with two retro-miRs (Figure 6A-B) in Liver Hepatocellular Carcinoma (LICH) and another with three retro-miRs with prognostic value in Stomach Adenocarcinoma (STAD), Figure 6C-D. In detail, Figure 6A shows the overall survival curve of two sets of LICH patients: patients with a higher expression of miR-4444 and miR-3654 present a significant (p-value < 0.0002) longer overall survival (e.g., overall survival of ∼1650 days for 50% of patients) than patients with a lower miR-4444 and miR-3654 expression (e.g., survival of ∼2600 days (60% more) for 50% of patients). For STAD patients, three retro-miRs (miR-622, miR-4788 and miR-4444) with higher expression were associated with worse overall survival. Patients with a higher expression of these retro-miRs had a lower overall survival (e.g., ∼850 days for 50% of patients) versus patients with a lower expression of these retro-miRs (e.g., survival of ∼1300 days for 50% of patients). Thus, these data also show a prognostic value of these retro-miRs in LIHC and STAD, and add a more layer of functionality (which need to be further explored) to these retrocopy derived miRNAs.

**Figure 6:**
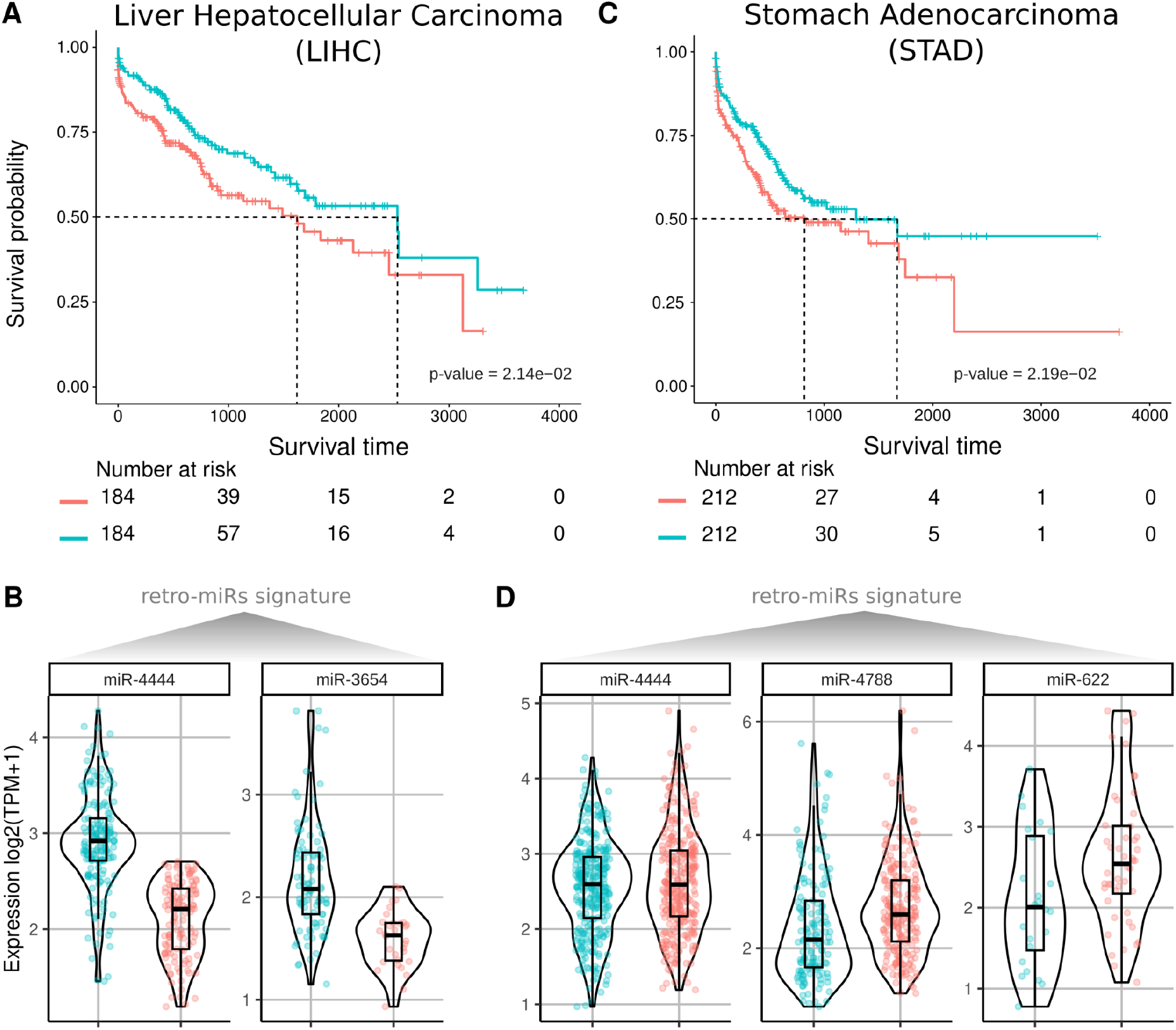
Prognostic value of retro-miRNA expression in liver hepatocellular carcinoma and stomach adenocarcinoma. **A)** Kaplan-Meier plot showing the overall survival differences (log-rank test) between two groups of patients with liver hepatocellular carcinoma, stratified based on expression levels of retro-miRs miR-4444-2 and miR-3654. **B)** Violin plot of the expression of miR-4444 and miR-3654 in the groups with better and worse overall survival probability presented in A. **C)** Kaplan-Meier plot showing the overall survival differences (log-rank test) between two groups of patients with stomach adenocarcinoma, stratified based on expression levels of retro-miRs miR-4444, miR-4788, and miR-622. **D)** Violin plot of the expression of miR-4444, miR-4788, and miR-622 in the groups with better and worse overall survival probability presented in C.

## DISCUSSION

MicroRNAs (miRNAs) are crucial regulators of gene expression and instrumental elements in species evolution. Understanding the mechanisms behind miRNA origination and their functions is essential for comprehending the evolution of increasingly complex organisms and disease development, including cancer. Retrocopies, gene copies generated by retroduplication of a mRNA molecule [18], are widespread in human and primate genomes and play crucial role in shaping the genomic landscape and driving evolutionary innovation [17,19]. In this study, we investigated retro-miRs, a class of miRNAs that emerged from the retroduplication of protein coding genes (mRNAs). We explored retro-miRs origination, conservation, expression, gene targets, and putative roles in cancer, including their potential to serve as tumor biomarkers and their association with overall cancer survival (prognostics).

MiRNAs are widespread through the animal kingdom [33]. The current miRBase release (22.1) presents 253 annotated (precursor) miRNAs in c. *Caenorhabditis elegan*, 882 mRNAs in chicken, 1,234 in mice, and 1,917 miRNAs in humans. Interestingly, some miRNAs are widely conserved (e. g., mir-100, conserved in eumetazoans), but mammals and especially primates present a high number of miRNAs, which is associated with the emergence of novel phenotypes in the evolution of these species [5]. Briefly, miRNAs act mainly by buffering (reducing the variance in) the expression level of their target genes. In the end, miRNAs ensure a stabilizing gene expression mode against endogenous and exogenous factors, maintaining invariant (novel) phenotypes [5,6]. The prerequisite for the origination of a novel mRNA is a transcribed genomic locus capable of producing a RNA fold recognizable by the miRNA processing machinery [1]. DNA duplication of pre-existing miRNAs, followed by their subfunctionalization or neofunctionalization, are the major sources of novel miRNAs genes [8]. However, miRNA origination from scratch (specially from intronic regions), from the duplication of non-coding RNAs (snoRNAs and tRNAs) and from retroduplicated elements (transposable elements) have been also reported [34]. Interestingly, Eric Devor [15] demonstrated that two primate-specific miRNAs originated in retroduplicated genes (processed pseudogenes or retrocopies) and suggested that retrocopies are good miRNA incubators [15]. Conversely, even with the burst in the genomics and bioinformatics, new investigations of miRNAs being originated in retrocopies have not been conducted. This work fills this gap in the literature, revealing 17 novel miRNAs originated by retroduplication of protein coding genes. Interestingly, these novel miRNAs are primate specific, with three of them being shared only between human and chimpanzee and two human specific retro-miRs, Figure 2. Surprisingly, some of these miRNAs (hsa-mir-572, hsa-mir-622) have been studied for years, but never detected as originated by a mRNA retroduplication. For example, the (retro-)mir-572 originated in RNPS1 retrocopy, RNPS1P1.

In order to function, a miRNA gene must be transcribed, folded, and processed, and it must have gene targets [1]. By using RNA sequencing data (and transcription start site definition) of tissues and cell lines, we confirmed that all retro-miRs (except mir-10527) are transcribed (Figure 3). Curiously, mir-10527 shows expression in cancer and in their normal counterpart tissues (Figure 5). As expected of a set of genes (miRNAs) presenting distinct functions and under the control of different promoter regions, the expression profile of retro-miRs ranges from retro-miRs expressed in few samples and tissues, to others being higher expressed and in more tissues (Figure 3). In a second step, we sought the retro-miRs gene targets. Notably, all retro-miRs have predicted and experimentally validated gene targets, except mir-10527, which lacks a validated target. Gene Ontology analysis of the biological processes associated with retro-miR targets revealed genes involved in fundamental cellular processes and functions, including cell differentiation and signaling, DNA replication, gene expression regulation, and neurotransmitter transport (Figure 4). Remarkably, two novel retro-miRs, mir-572 and mir-4788, have gene targets that are primarily involved in brain function, such as nervous system development, regulation of neurotransmitter levels, neurotransmitter transport, and synapse. Therefore, since miRNAs provide robustness to their target genes’ regulatory networks, helping to maintain novel traits [5], these novel and primate-specific retro-miRs are promising candidates for further exploration in the context of key genes (miRNAs) to explain the evolution of the brain in humans and other primates.

Additionally, we found that retro-miRs target important cancer-related genes (Figure 4D), which prompted us to investigate their expression in cancer. First, we confirmed that retro-miRs are highly expressed in several cancer types (Figure 5). Interestingly, three retro-miRs (mir-572, mir-622, and mir-492) have been more extensively studied in cancer. Studies have shown that miR-572 can act as both a tumor suppressor and an oncogene, depending on the tissue and tumor type. Consistent with our findings (Figure 5), miR-572 has been shown to have low expression levels (potentially acting as a tumor suppressor) in breast [35] and colorectal tumors [36]. On the other hand, the oncogenic role of mir-572 has been confirmed in lung tumors [37], hepatocellular carcinoma [38], and kidney cancer [39], which is also consistent with our results (Figure 5). However, the importance of this retro-miR in other tumors (Figure 5) remains underexplored. For miR-622, the literature also reports that this retro-miR can act as both a tumor suppressor or an onco-miR in several cancers, such as breast cancer, glioma, CRC, hepatocellular carcinoma (HCC), lung cancer, gastric cancer (GC), melanoma, ovarian carcinoma, prostatic cancer, and pancreatic cancer [40], in agreement with our expression results (Figure 5). Like the other two retro-miRs, the literature reports that miR-492 acts as both a tumor suppressor and an oncogene. In agreement with our expression data, this retro-miR is upregulated in metastatic hepatoblastoma [41], lung cancer [42], and stomach cancer [43]. However, for the other retro-miRs (Figure 5), few or no investigations have been performed in tumor tissues.

In a final investigation, we also found that two sets of retro-miRs have their expression (signature) associated with patient overall survival of two cancer types, Liver Hepatocarcinoma (LIHC) and Stomach Adenocarcinoma (STAD), Figure 6. These results should be further investigated.

## CONCLUSIONS

In conclusion, we have conducted the most comprehensive study to date on miRNAs that are derived from retroduplicated protein-coding genes in the human genome. Our findings demonstrate that these retro-miRs are expressed, are located in a conserved environment, have regulatory targets in fundamental cellular processes - including those in the nervous system - and exhibit differential expression and functional roles in various cancer types. Overall, our study sheds light on the role of mRNA retrotransposition in generating driver genetic novelties, such as miRNAs, and highlights the potential for future investigations into the function of retrocopies in humans and other organisms.

## METHODS

### Identification of retro-miRs derived from retrocopies

To identify the retrocopy derived miRNAs (retro-miRs) we followed these steps: i) We used mirBase (http://www.mirbase.org/) as an annotation source for miRNAs characteristics, such as their genomic position and gene name. ii) Our set of retrocopied genes was obtained from the RCPedia database [26], which contains several information about retrocopies (e.g., genomic location, sequences, expression and conservation) and their parental genes; In order to convert the retrocopies’ genomic coordinates from hg19 to hg38 we used the LiftOver online tool (https://genome.ucsc.edu/cgi-bin/hgLiftOver). Subsequently, we selected the miRNAs from miRBase whose genomic coordinates overlapped with those of retrocopies using BedTools v2.26.0 (with default parameters and a minimum overlap of 5 bp) [44]. We manually removed miRNAs that originated from an insertion of a Mobile Element inside the retrocopies. To assess the co-localization (exonic, intronic and intergenic) of the miRNAs and protein coding genes we utilized the miRIAD database [24,25]. Additionally, we selected annotated exonic miRNAs of coding genes that have retrocopies and the retro-miRs already identified. We aligned the mature miRNA sequence from these candidates against the respective retrocopies of the coding gene using Clustal Omega available at EMBL (https://www.ebi.ac.uk/Tools/msa/clustalo/). To evaluate the formation of secondary structures we used RNAFold version 2.4.17 [45]. When the secondary structure is formed, the candidates are considered retro-miRs.

### DNA-mediated Duplication

To identify duplicated genes, we used the transcripts annotated by GENCODE v32. We used tblastn 2.7.1+ [46] with default parameters to align the amino acid sequences of all transcripts against the nucleotide sequences of all transcripts. We considered that a gene has a DNA-mediated duplication if it has at least one alignment besides their own nucleotide transcript that has a score, that divided by the score of their original nucleotide transcript, is greater or equal to 0.7. Additionally, we searched for miRNAs in these duplicated genes and for duplications of these miRNAs in the duplicated genes using the miRNAs genomic coordinates provided by miRBase.

### Classification of retro-miRs

To classify retro-miRs into Retroposed (R-retro-miRs), Exon Junction EJ-retro-miRs and Novel retro-miRs (N-retro-miRs), we aligned the sequences of the retro-miRs against the reference genome (GRCh38) using the BLAT tool (https://genome.ucsc.edu). For each retro-miR, we investigated the pattern of alignment against the parental gene of the retrocopy. Based on this, 3 major patterns emerged: (I) retro-miRs that are exact copies of the exonic miRNA of the parental sequence (R-retro-miRs); (II) retro-miRs originated from exon-exon junction (EJ-retro-miRs); (III) retro-miRs originated from mutations (>1 bp) in relation to the parental gene in the mature sequence (N-retro-miRs).

### Conservation of retro-miRs among primates

The conservation of retro-miRs among primate genomes was done through MultiZAling available in the Genome Browser (https://genome.ucsc.edu/index.html). We compared the sequences of miRNAs annotated in humans with those of the genomes of Chimpanzee (Pan_tro 3.0), Gorilla (GSMRT3), Orangutan (WUGSC 2.0.2), Rhesus (BCM Mmul_8.0.1) and Marmoset (WUGSC 3.2). We considered a retro-miR to be conserved if the sequence identity with the homologous sequences of these genomes was greater than 80%.

### dN/dS Analysis

To calculate the nonsynonymous to synonymous substitution rate ratio (dN/dS), we extracted the nucleotide sequences of the retrocopies from hg38, and we used ORFfinder [47] to extract the longest predicted aminoacid (aa) sequence for each of them. In the cases of retrocopies that are human specific (HNRNPA3P6 and RPS27AP5), we used BLAT [48] to align the predicted aa sequence against the human genome (version GRCh38/hg38) and retrieve the best match on the parental gene nucleotide sequence (paralog). For retrocopies with a primate ortholog (RPS27AP16, PTMAP2, PTMAP9, PTMAP4, RP11-371A22.1, RNPS1P1, PTMAP8, RP11-529H20.3, EEF1GP5, KRT18P27, TATDN2P2, HMGB3P13, RCC2P3, KRT19P2, PABPC1P4), we align the human predicted aa sequence against the genome of chimpanzees (panTro6) with BLAT and retrieve the best match on the retrocopy ortholog. For both the paralog and ortholog, we use ORFfinder to find the longest predicted aa sequence that is equivalent to the aa sequence of the human retrocopy. We used ClustalW [49] to align the aa sequence of the retrocopy and their correspondent in the paralog or ortholog. PAL2NAL (codeml; CodonFreq = 2, model = 0, Nsites = 0, fix_omega = 0, omega = 0.4) [50] was used to calculate the synonymous (dS) and non-synonymous (dN) substitution rates. p-values of dN/dS were determined by comparison to a model assuming neutral evolution (fix_omega = 1, omega = 1 for paralogs and omega = 0.5 for orthologs) with a Likelihood-ratio-test [LRT], p<0.05, chi-sq. distribution. blastn [47] was used to calculate the identity between the nucleotide sequence equivalent of the CDS.

### Retro-miRs expression and transcription start site definition in normal and cancer tissues

The expression of these retro-miRs in normal tissues were analyzed using pre-processed data from the FANTOM phase 5 database [29,30]. The extracted data were related to the expression of retro-miRs by miRNA-Seq in samples of different cell types (Table 1S). We search for a transcription start site (TSS) on the expected transcription orientation (based on the miRNA direction) up to a distance of 4k base pairs of the 5’ end of the retro-miR. We also used miRNAs expression pre-processed (short RNA sequencing methodology) data from The Cancer Genome Atlas (TCGA) and microRNA Tissue Expression Database (miTED). We selected the TCGA data as the main data for further investigations and selected miTED expression data for miRNAs that were not initially expressed in the TCGA data. In both datasets we normalized this data by transcript per million (TPM). Expression is quantified only for the mature sequence of the retro-miRs.

### Investigating the prognostic value of retro-miRs in 10 cancer types

To investigate the prognostic value of retro-miRs in cancer, expression (short RNA-sequencing methodology) and patient’s survival information were extracted from TCGA for the following tumor types: Breast cancer (BRCA), Colon adenocarcinoma (COAD), Kidney renal clear cell carcinoma (KIRC), Liver hepatocellular carcinoma (LIHC), Lung squamous cell carcinoma (LUSC), Lung adenocarcinoma (LUAD), Pancreatic adenocarcinoma (PAAD), Prostate adenocarcinoma (PRAD), Stomach adenocarcinoma (STAD), Thyroid carcinoma (THCA). Only retro-miRs expressed (TPM > 0) were considered in the analysis. The association between retro-miRs expression signature (combined expression of two or more retro-miRs) and patients overall survival was assessed using Reboot (dos Santos et al., 2021) with default parameters.

### Finding retro-miRs’ target genes

To find out the target genes of retro-miRs, we ran TargetScan 7.0 with the set of 3′ UTR sequences of protein-coding genes provided by the Targetscan [27]. We selected only transcripts that had 8mer-1a or 7mer-m8 site types or if these genes were represented in miRTarBase [28]. Enrichment analyses of target genes in Gene ontology (Biological Processes) and KEGG pathways were performed using ShinyGO v0.77 (http://bioinformatics.sdstate.edu/go/). Redundant Gene Ontology terms were further removed using REVIGO (http://revigo.irb.hr/). Only GO processes and KEGG pathways with fold enrichment > 1.5 and FDR<0.05 were considered as significant.

## DECLARATIONS

### Ethics approval and consent to participate

Not applicable.

### Consent for publication

Not applicable.

### Availability of data and materials

The datasets used and/or analyzed during the current study are available from the corresponding author on reasonable request.

### Authors’ contributions

R.L.M. and P.A.F.G. conceived the study. R.L.M., H.B.C., and G.D.A.G. performed the analyses. G.G., M.D.V., and L.C.H. contributed to the analyses. R.L.M., H.B.C., G.D.A.G., G.G., M.D.V., L.C.H., and P.A.F.G. discussed the results. P.A.F.G. coordinated the study and prepared the manuscript, with inputs from all authors. All authors reviewed and approved the final manuscript.

## Acknowledgements

We would like to thank all members of the Galante lab for their helpful discussions and feedback on the manuscript.

