## Supplementary Figures for "Retro-miRs: Novel and functional miRNAs originated from mRNA retrotransposition"

### SUPPLEMENTARY DATA

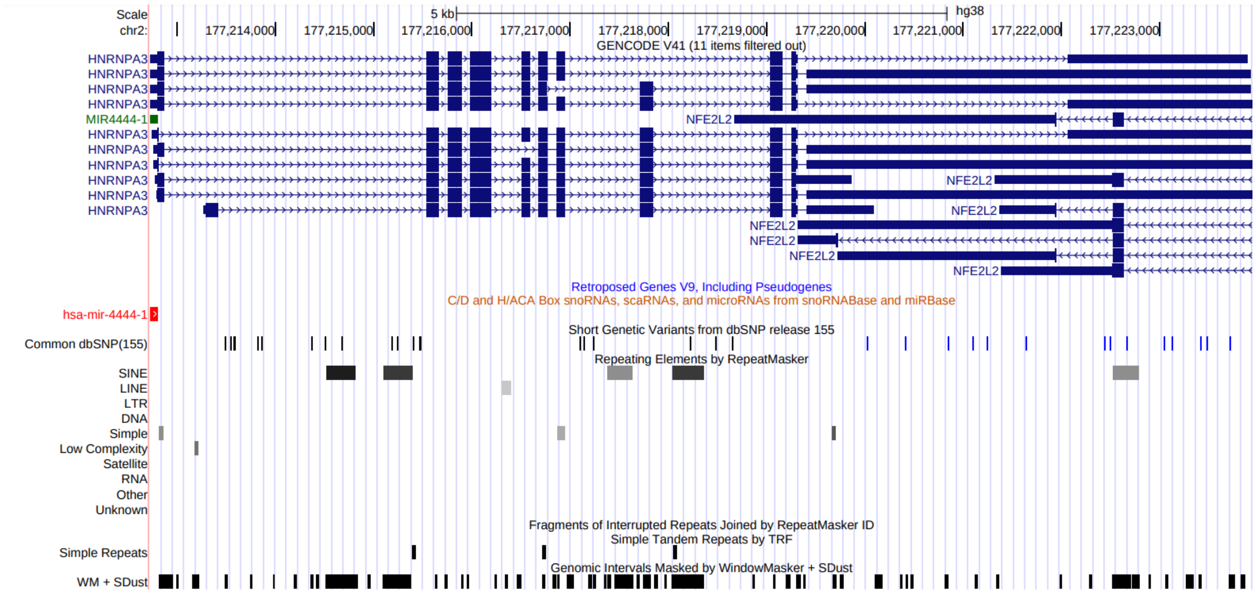

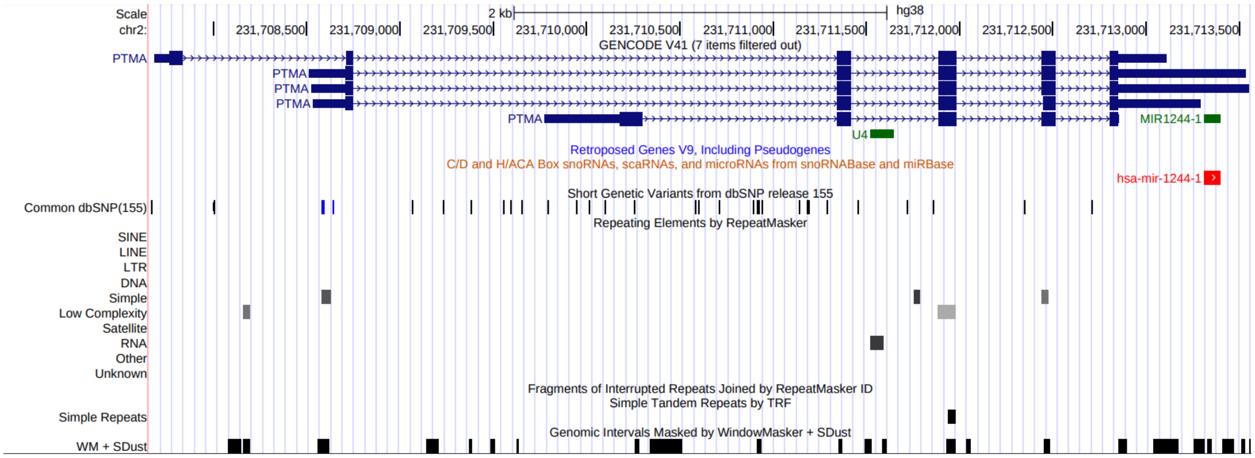


**Figure S1**. Exonic miRNAs already present in their parental gene sequences

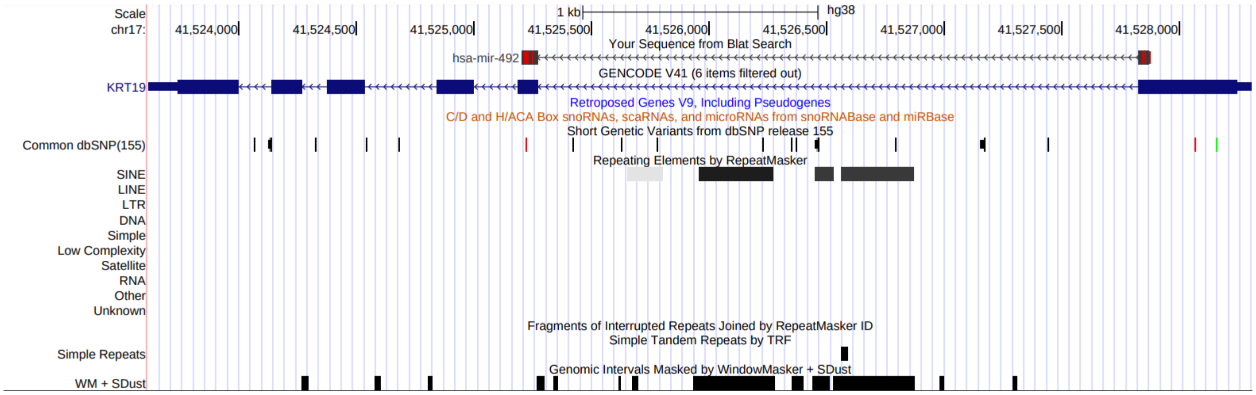


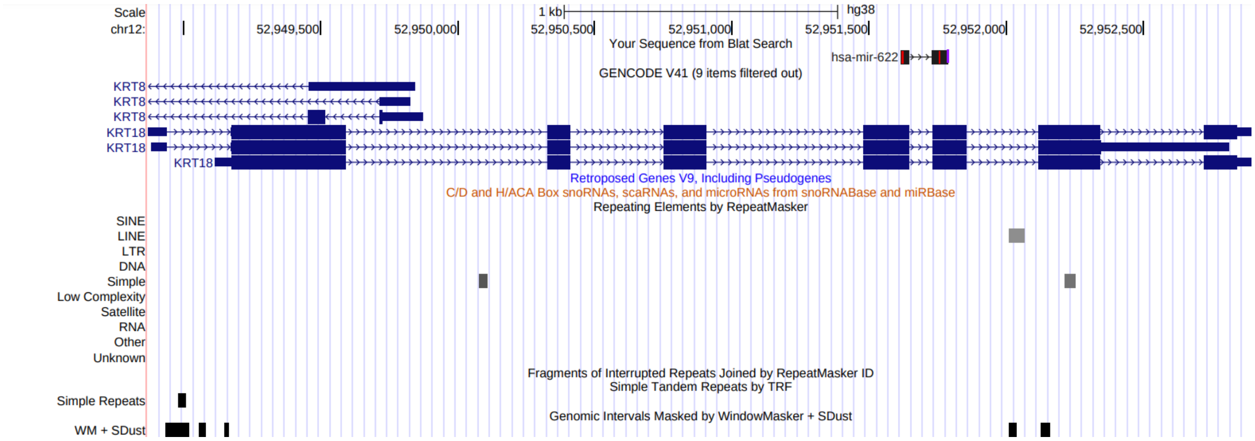

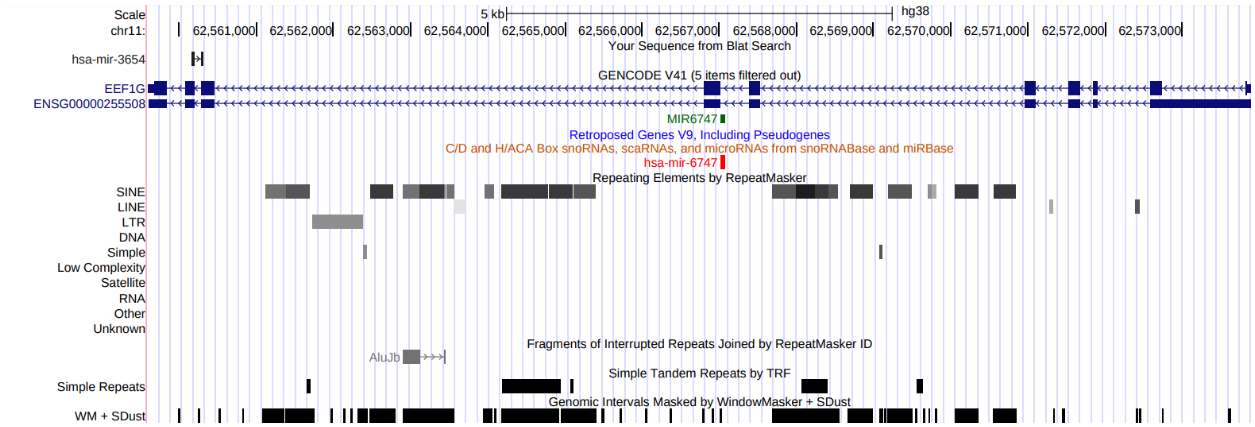
Scale
chr11:
5 kb
hg38
62,561,000
62,562,000
62,563,000
62,564,000
62,565,000
62,566,000
62,567,000
62,568,000
62,569,000
62,570,000
62,571,000
62,572,000
62,573,000
Your Sequence from Blat Search
hsa-mir-3654
GENCODE V41 (5 items filtered out)
EEF1G
ENSG00000255508
MIR6747
Retroposed Genes V9, Including Pseudogenes
C/D and H/ACA Box snoRNAs, scaRNAs, and microRNAs from snoRNABase and miRBase
hsa-mir-6747
Repeating Elements by RepeatMasker
SINE
LINE
LTR
DNA
Simple
Low Complexity
Satellite
RNA
Other
Unknown
Fragments of Interrupted Repeats Joined by RepeatMasker ID
AluJb
Simple Tandem Repeats by TRF
Simple Repeats
Genomic Intervals Masked by WindowMasker + SDust
WM + SDust

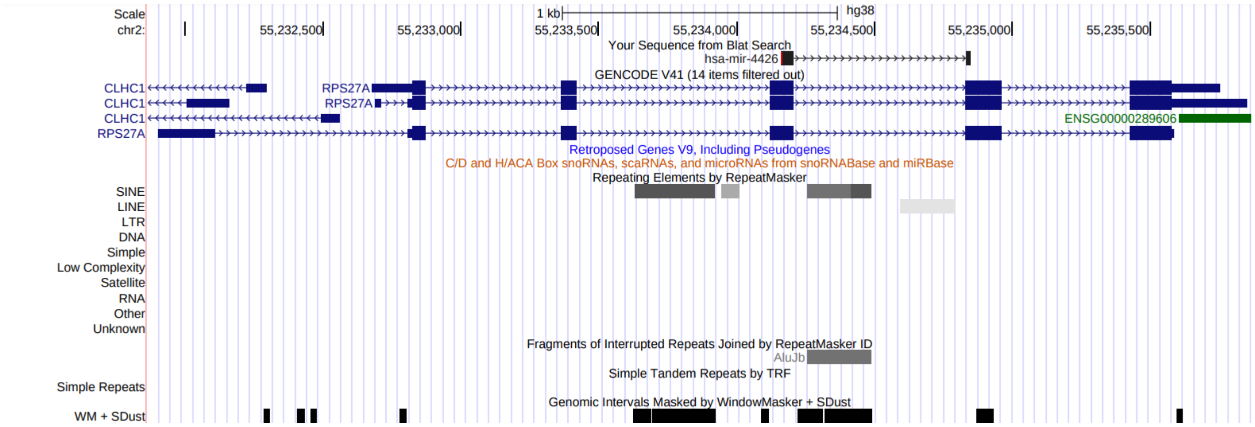
Scale
chr2:
1 kb
hg38
55,232,500
55,233,000
55,233,500
55,234,000
55,234,500
55,235,000
55,235,500
Your Sequence from Blat Search
hsa-mir-4426
GENCODE V41 (14 items filtered out)
CLHC1
RPS27A
CLHC1
RPS27A
CLHC1
RPS27A
ENSG00000289606
Retroposed Genes V9, Including Pseudogenes
C/D and H/ACA Box snoRNAs, scaRNAs, and microRNAs from snoRNABase and miRBase
Repeating Elements by RepeatMasker
SINE
LINE
LTR
DNA
Simple
Low Complexity
Satellite
RNA
Other
Unknown
Fragments of Interrupted Repeats Joined by RepeatMasker ID
AluJb
Simple Tandem Repeats by TRF
Simple Repeats
Genomic Intervals Masked by WindowMasker + SDust
WM + SDust

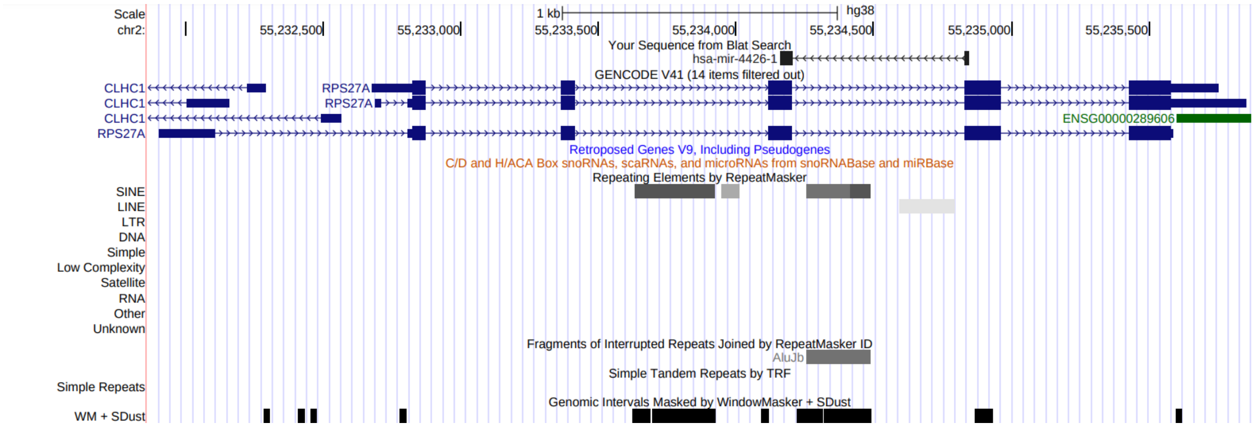
Scale
chr2:
1 kb
hg38
55,232,500
55,233,000
55,233,500
55,234,000
55,234,500
55,235,000
55,235,500
Your Sequence from Blat Search
hsa-mir-4426-1
GENCODE V41 (14 items filtered out)
CLHC1
RPS27A
CLHC1
RPS27A
CLHC1
RPS27A
ENSG00000289606
Retroposed Genes V9, Including Pseudogenes
C/D and H/ACA Box snoRNAs, scaRNAs, and microRNAs from snoRNABase and miRBase
Repeating Elements by RepeatMasker
SINE
LINE
LTR
DNA
Simple
Low Complexity
Satellite
RNA
Other
Unknown
Fragments of Interrupted Repeats Joined by RepeatMasker ID
AluJb
Simple Tandem Repeats by TRF
Simple Repeats
Genomic Intervals Masked by WindowMasker + SDust
WM + SDust

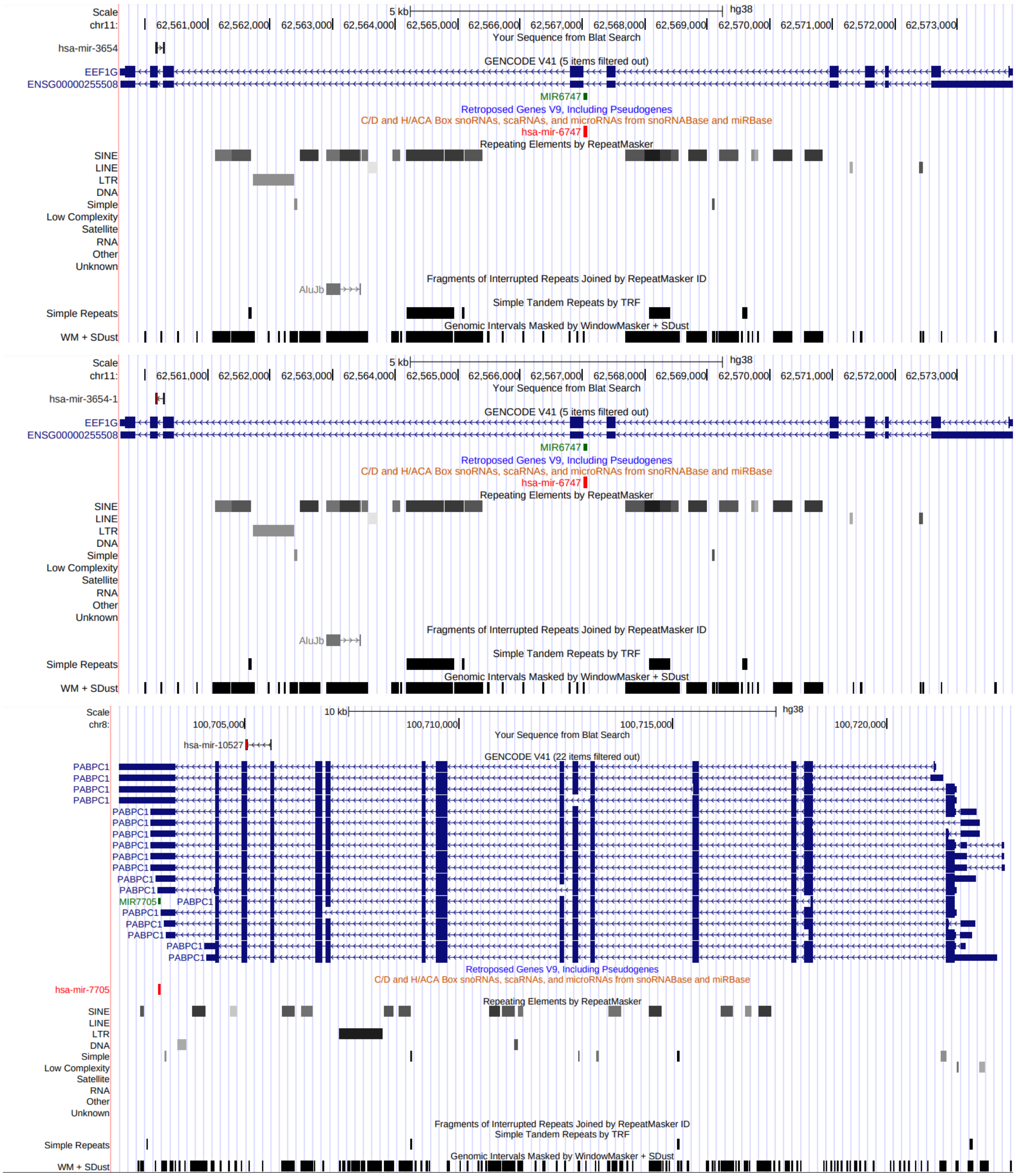


**Figure S2.** Retro-miRs spanning two exons.

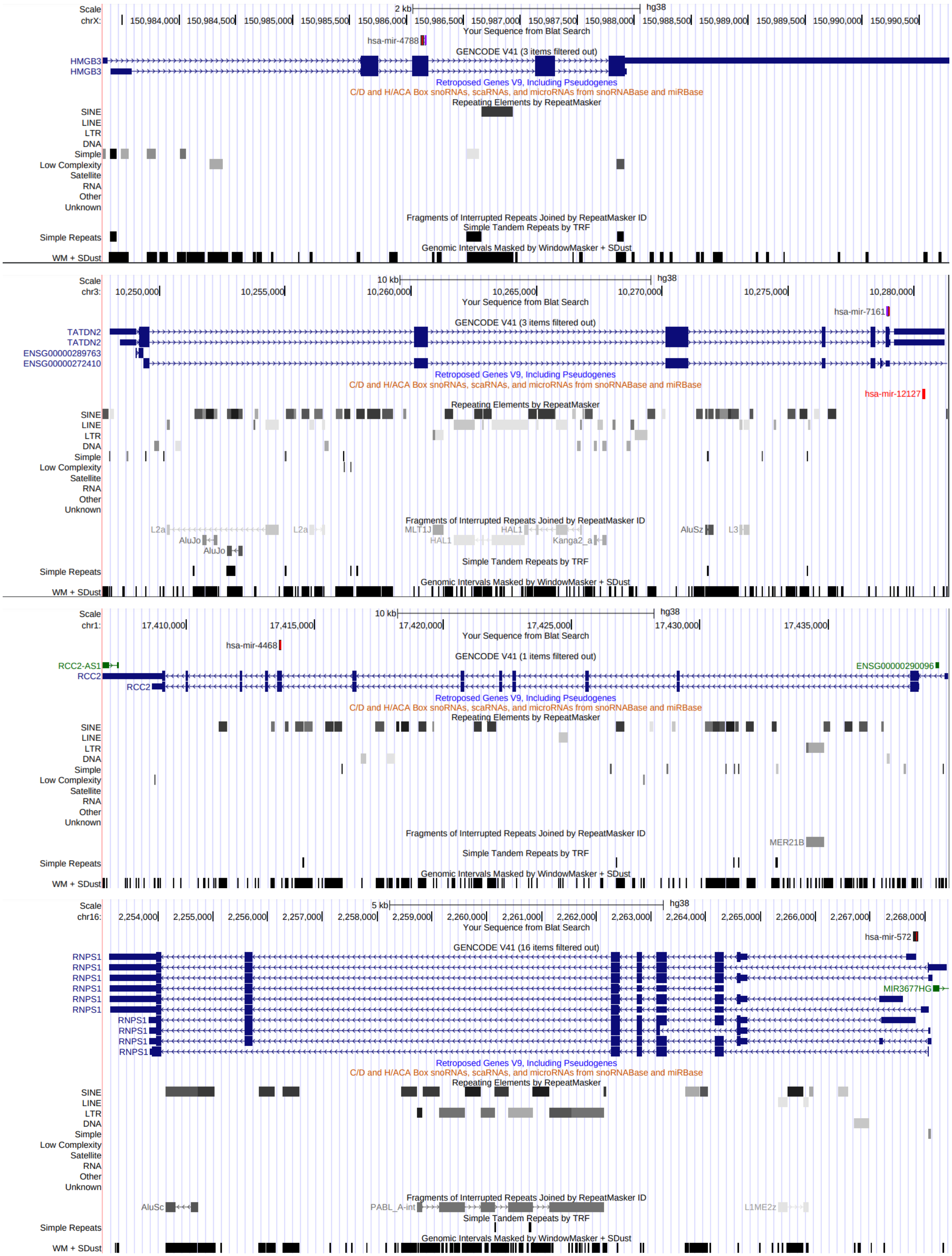


Figure S3. Parental genes without exonic miRNA.

| | |
|---|---|
| RNPS1 | chr16:2267786-2267880 |

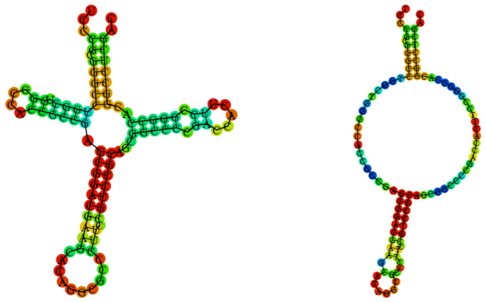

| | |
|---|---|
| mir-572 | chr4:11368827-11368921 |

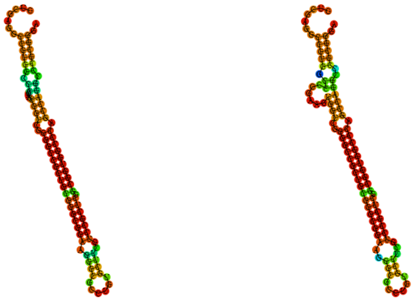

| | |
|---|---|
| TATDN2 | chr3:10278942-10279024 |

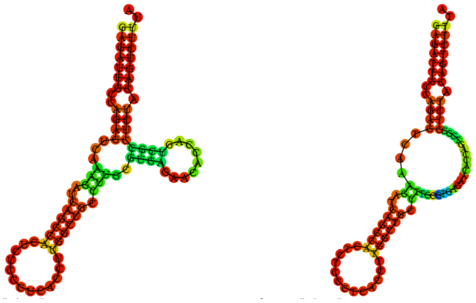

| | |
|---|---|
| mir-7161 | chr6:158609707-158609790 |

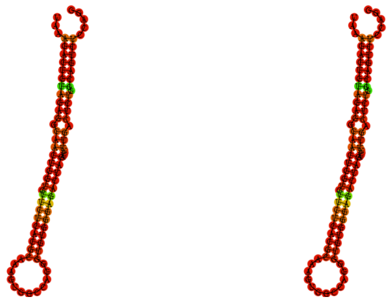

| | |
|---|---|
| HMGB3 | chrX:150986122-150986170 |

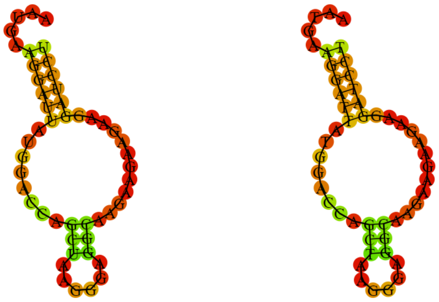

| | |
|---|---|
| mir-4788 | chr3:134437827-134437906 |

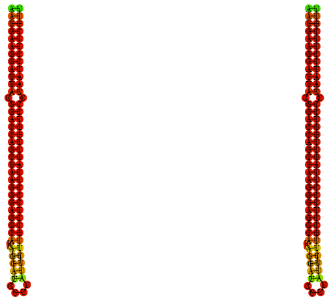

| | |
|---|---|
| RCC2 | chr1:17413631-17413694 |

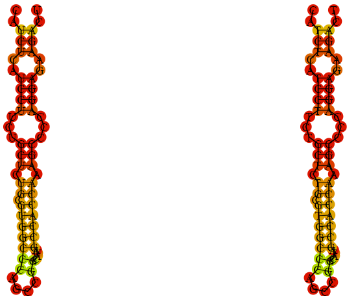

| | |
|---|---|
| mir-4468 | chr7:138123758-138123821 |

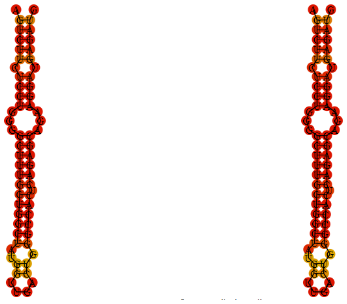

Figure S4. Regions with and without evidence of a stem loops.

miR-10527

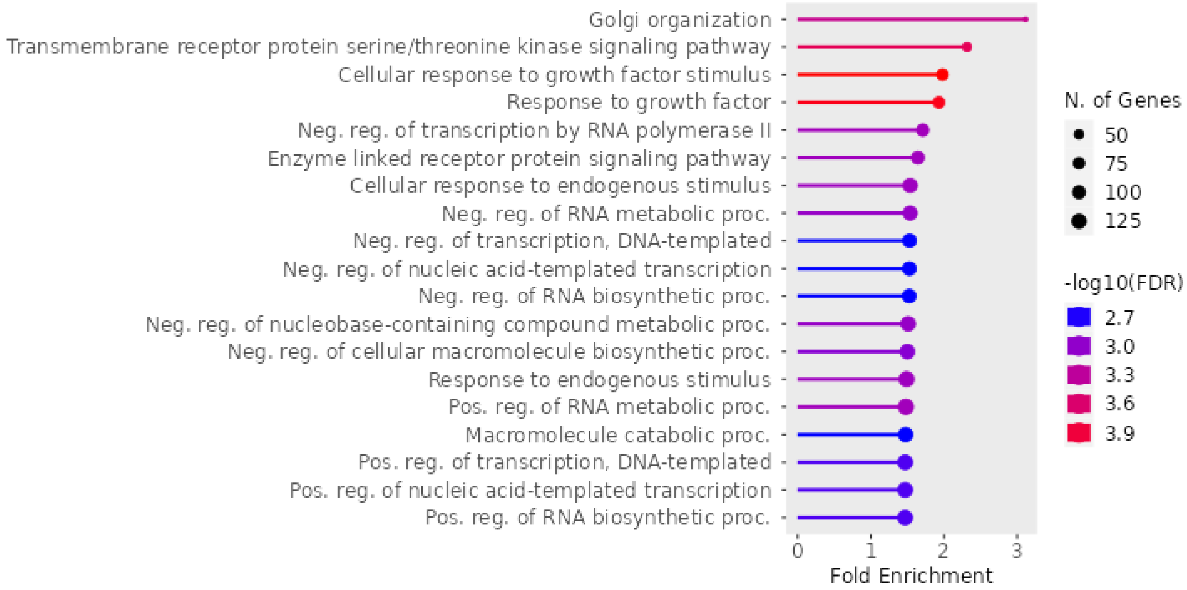


miR-1244

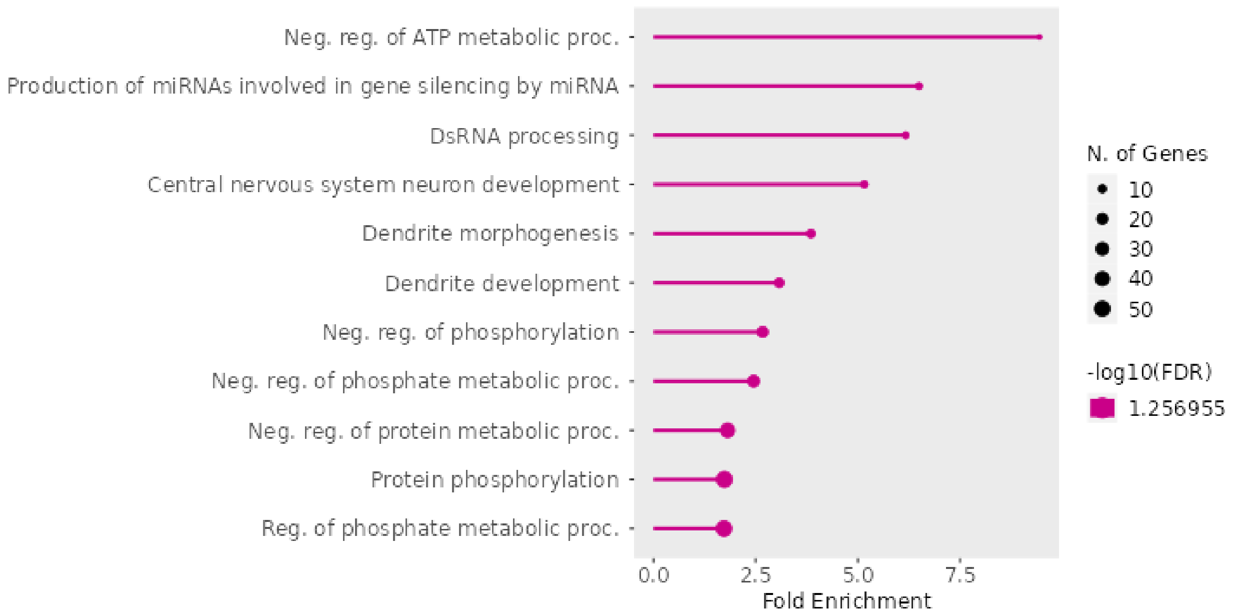


miR-3654

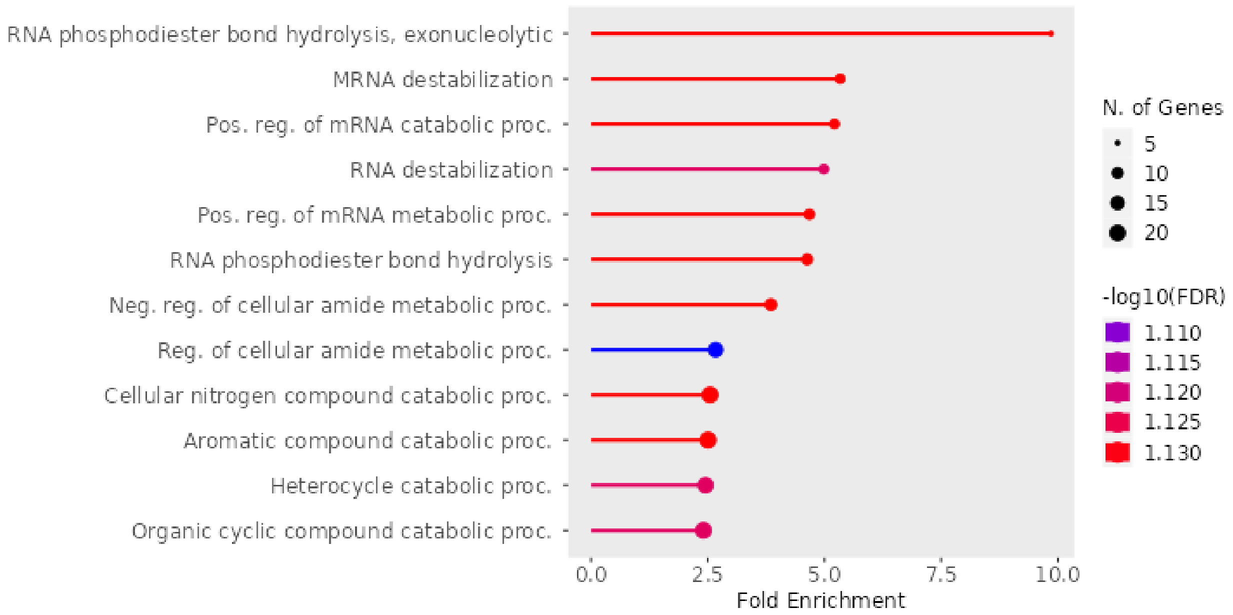


miR-4426

None

miR-4444

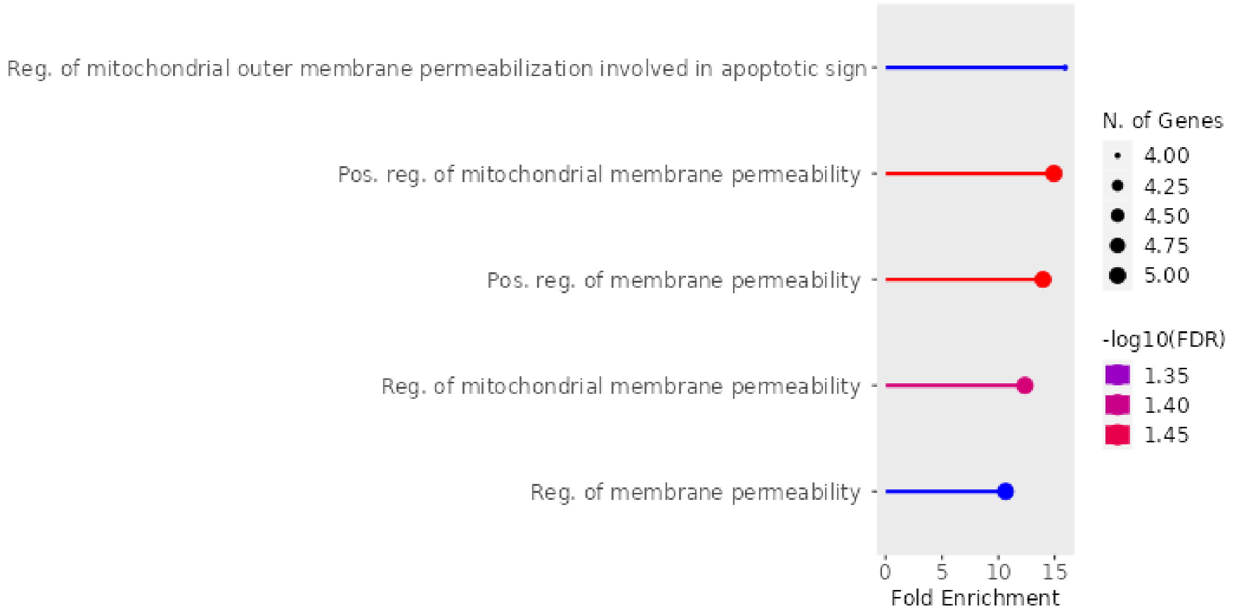


miR-4468

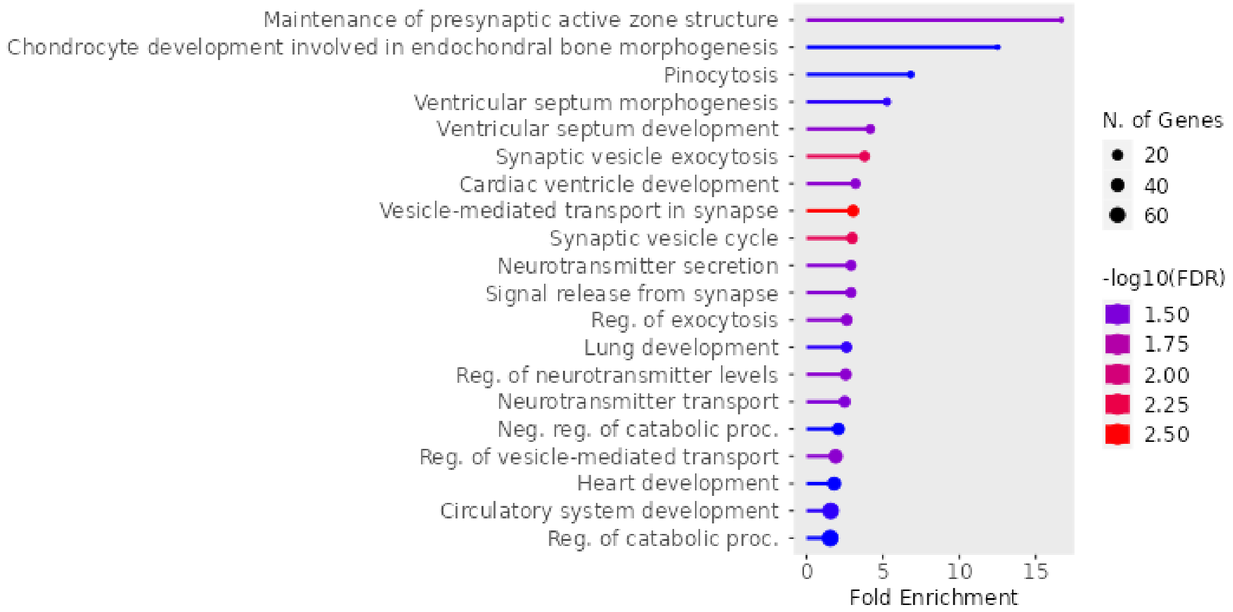


miR-4788

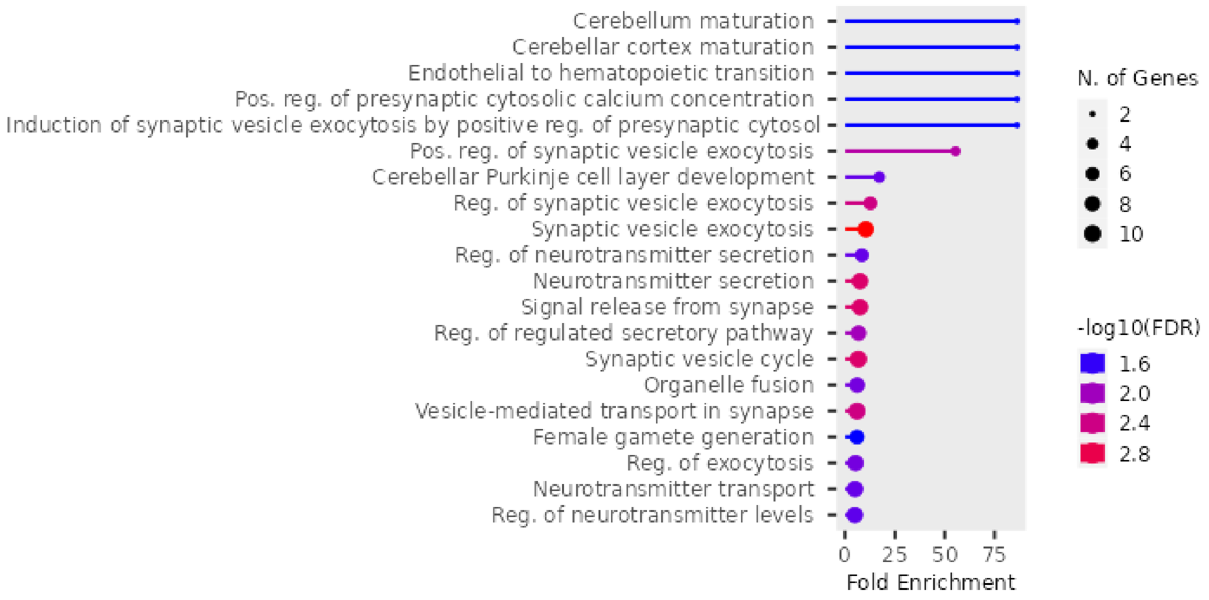


miR-492

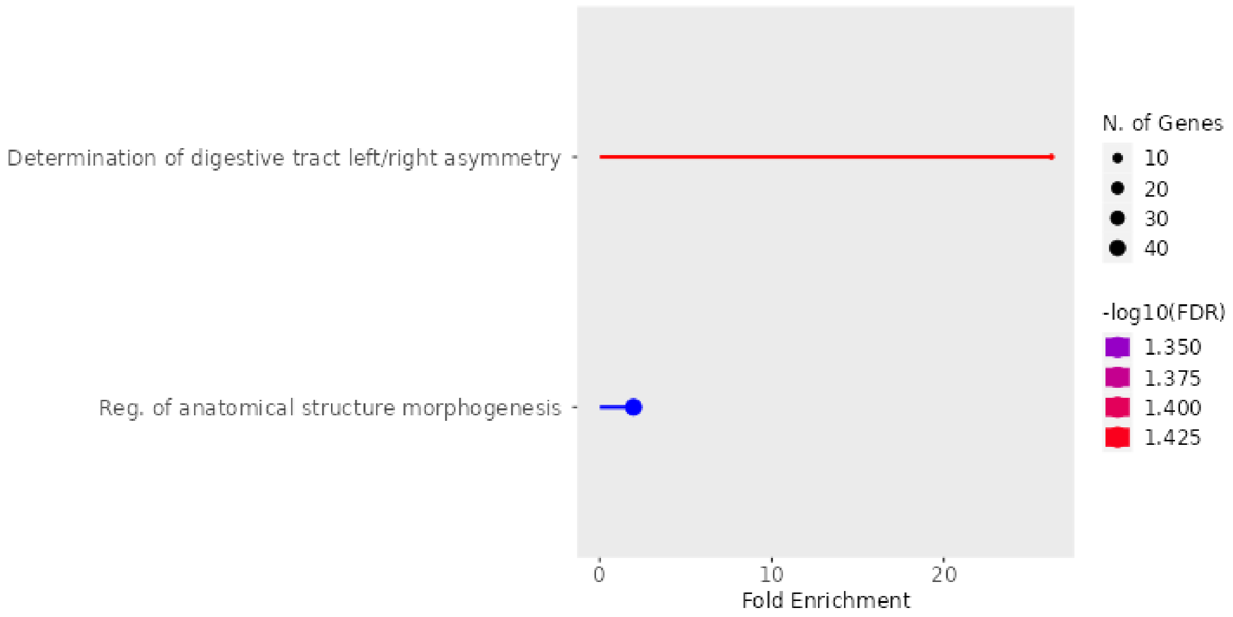


miR-7161-5p

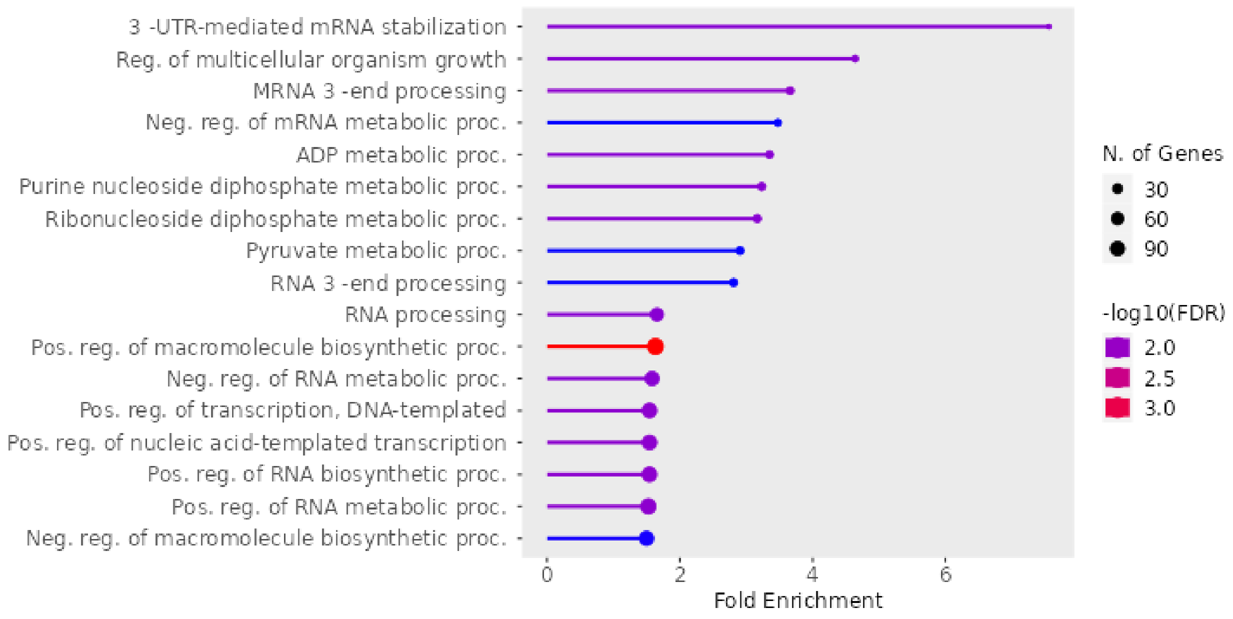


miR-7161-3p
None

miR-572

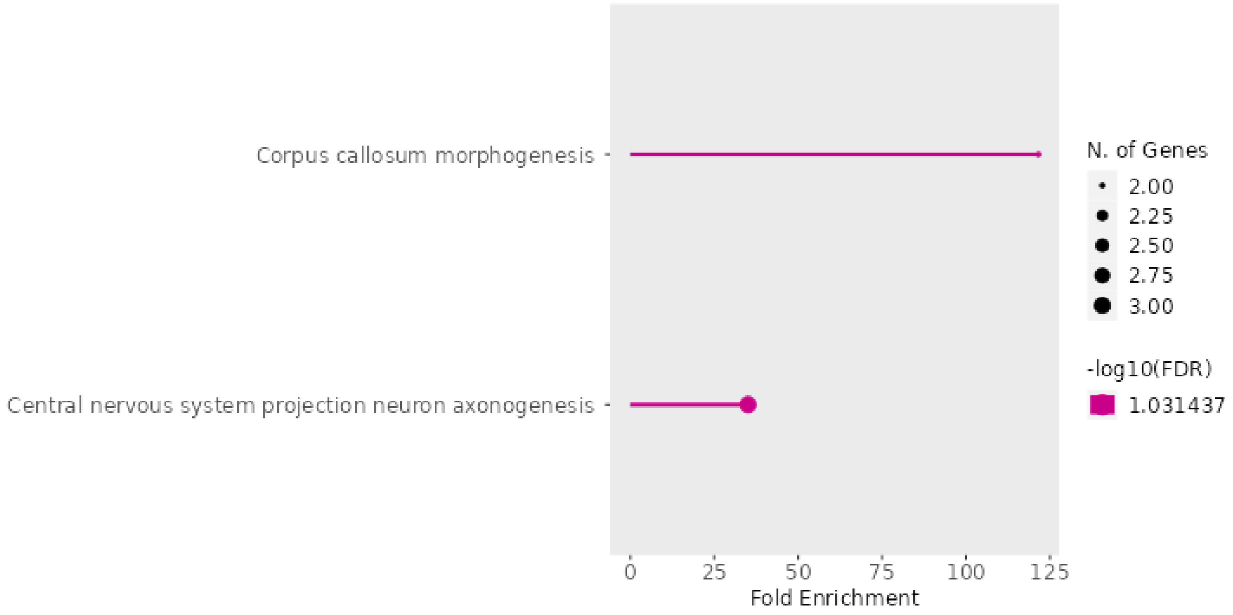


miR-622

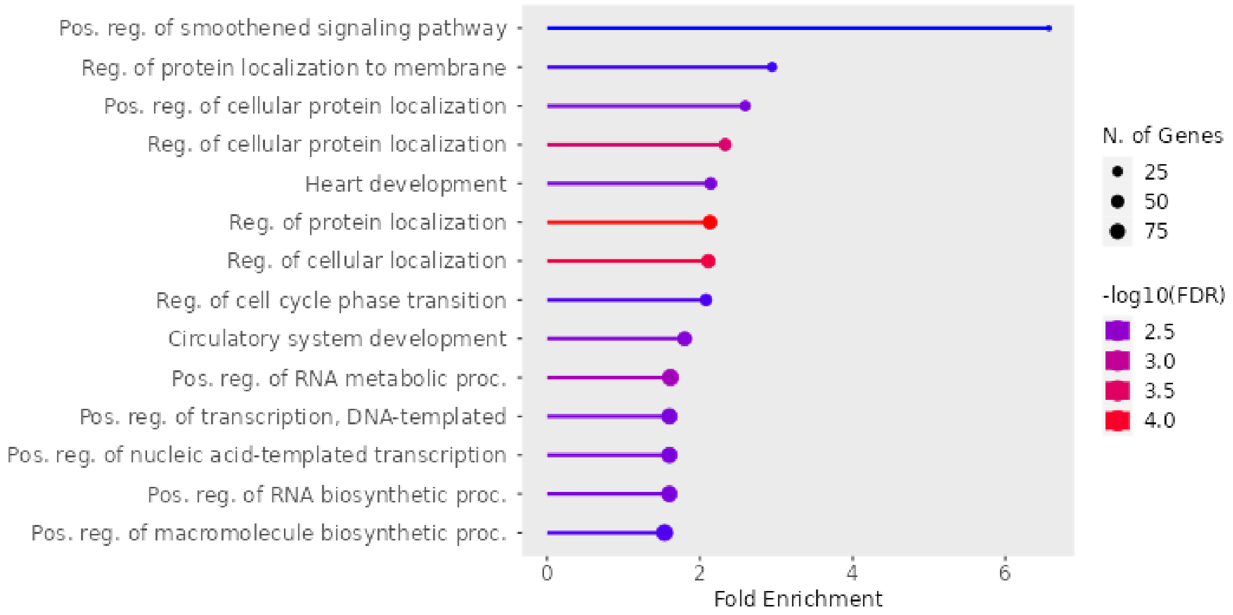


Figure S5. Gene ontology analyses of retro-miRs targets.
